# MOLECULAR VIDEOGAMING: SUPER-RESOLVED TRAJECTORY-BASED NANOCLUSTERING ANALYSIS USING SPATIO-TEMPORAL INDEXING

**DOI:** 10.1101/2021.09.08.459552

**Authors:** Tristan P. Wallis, Anmin Jiang, Huiyi Hou, Rachel S. Gormal, Nela Durisic, Giuseppe Balistreri, Merja Joensuu, Frédéric A. Meunier

## Abstract

Single-molecule localization microscopy (SMLM) techniques are emerging as vital tools to unravel the nanoscale world of living cells. However, current analysis methods primarily focus on defining spatial nanoclusters based on detection density, but neglect important temporal information such as cluster lifetime and recurrence in “hotspots” on the plasma membrane. Spatial indexing is widely used in videogames to effectively detect interactions between moving geometric objects. Here, we use the R-tree spatial indexing algorithm to perform SMLM data analysis and determine whether the bounding boxes of individual molecular trajectories overlap, as a measure of their potential membership in nanoclusters. Extending the spatial indexing into the time dimension allows unique resolution of spatial nanoclusters into multiple spatiotemporal clusters. We have validated this approach using synthetic and SMLM-derived data. Quantitative characterization of recurring nanoclusters allowed us to demonstrate that both syntaxin1a and Munc18-1 molecules transiently cluster in hotspots on the neurosecretory plasma membrane, offering unprecedented insights into the dynamics of these protein which are essential to neuronal communication. This new analytical tool, named Nanoscale Spatiotemporal Indexing Clustering (NASTIC), has been implemented as a free and open-source Python graphic user interface.

## INTRODUCTION

In recent years, great advances have been made in our understanding of cellular molecular dynamics, through the emergence of super-resolution microscopy, and in particular, a suite of technologies collectively referred to as single-molecule localization microscopy (SMLM)^1^. When applied in live cells, this approach allows the detection and tracking of individual molecules to be determined at nanometre scale, far below the diffraction limit of visible light, thereby opening the way to understanding the nanoscale world of living cells^2^. Single-molecule tracking at the level of the plasma membrane has revealed that membrane-associated proteins can congregate into functional assemblies called nanoclusters. Functional nanoclustering can bring receptors and effectors/substrates into close proximity at discrete regions of the plasma membrane. SMLM has allowed the characterization of molecular nanoclustering of synaptic receptors and their functions^3,4^. The formation of these nanoclusters can be further studied in live cells, allowing dynamic characterization of single receptors and their effectors confined into these discrete areas of the plasma membrane^5-13^. These studies seek to define metrics such as mobility, nanocluster size, lifetime, molecular membership and density, and establish how they correlate with changing experimental conditions. The key to such investigations is a robust analysis pipeline for determining which of the hundreds/thousands of molecular detections (Fig. 1A) acquired in a typical single-particle tracking experiment are confined into nanoclusters. To date, algorithms for nanocluster determination have largely relied on one of two fundamental principles: density/proximity assays such as DBSCAN (density-based spatial clustering of applications with noise)^14^ which determines clustering based on whether the number of molecular detections within a determined radius exceed a user-defined threshold (Fig. 1B), and segmentation-based assays such as Voronoï tessellation^15,16^, which define minimum tiles around each molecular detection and assign clustered detections based on a user-defined tile area threshold (Fig. 1C). More recently, computer vision algorithms have also been used to determine clustering based on algorithmic identification of features in SMLM data^17^. However, these techniques have been mostly applied for fixed cell data, and they are therefore lacking the temporal analysis which is useful when considering single particle tracking data sets.

**Figure 1.**
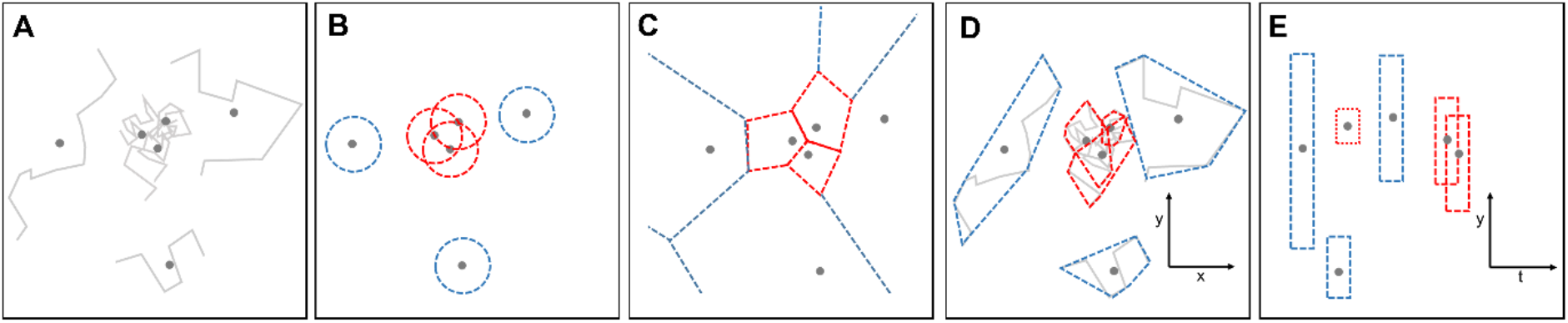
Schematic representation of clustering algorithms as applied to molecular trajectories. **(A)** Molecular trajectory data, with each trajectory’s spatial centroid indicated with a dot. (**B)** DBSCAN. Multiple molecular centroids present within a defined radius (red circles) are considered clustered. The most effective radius (**ε**) and minimum number of centroids within it (MinPts) are determined empirically. (**C)** Voronoï tessellation. Tiles are drawn around each centroid such that the distance from any point within the tile is closer to its centroid than to any other centroid. Molecular centroids with tile areas less than an empirically determined threshold (red) are considered clustered. (**D)** Spatial indexing. Clustered molecules are determined by overlapping 2D bounding regions (red) defining the spatial extent of each molecular trajectory. (**E)** Spatiotemporal indexing. This panel represents the data in panel D rotated 90° around its y axis to highlight the temporal component of each centroid. Each trajectory bounding region is assigned a user-defined “thickness” in the time dimension. Overlapping 3D bounding regions represent spatiotemporally clustered molecules.

In this study, we propose an alternative approach to examining molecular clustering in live-cell SMLM data, which is based upon two primary assertions: 1) in live-cell data the molecular trajectory is the indivisible unit for each tracked molecule, not the individual molecular detections comprising the trajectory, and 2) trajectories which spatially and temporally overlap with other trajectories are more likely to represent molecules that are confined into functional nanoclusters. We therefore consider that the ability to simultaneously derive information about the spatial and temporal interactions of molecular trajectories can provide valuable insights into functional protein interactions on the cell membrane.

Accordingly, we investigated the use of bounding regions encompassing the extent of each molecular trajectory over the lifetime of its observation. Overlapping bounding regions represent an increased likelihood that their underlying molecules constitute members of a nanomolecular spatial cluster (Fig. 1D). Determining whether geometric shapes overlap is potentially computationally complex, particularly if there are large numbers of these shapes. This can be overcome using spatial indexing, which allows rapid querying of a spatial database of the shapes’ rectangular bounding boxes. Over the last few decades a number of approaches to spatial indexing such as Quad-tree^18^ and R-tree^19^ have been implemented, variations of which have found wide use in mapping and database management, as well as in videogames, where they allow accurate spatiotemporal detection of interactions between objects such as bullets and combatants^20^. The need to maintain high frame rates means that these routines have been highly optimized. Considering that spatiotemporal interaction is highly relevant, particularly in defining nanoclustering of molecules of interest, we used the R-tree spatial indexing algorithm to determine the overlap of the bounding regions of hundreds/thousands of trajectories. R-tree databases are k-dimensional, so in addition to the x/y spatial extent, we added an additional time extent (trajectory detection time ± user-defined time window). This approach allowed determination of trajectory overlap in space and time (Fig. 1E), and introduced a temporal component into the clustering metrics, allowing us to determine cluster lifetime, rate of cluster formation, and “hotspot” in discrete areas of the plasma membrane where detections occur intermittently. Using both synthetic and experimentally derived SMLM data, we compared <u>na</u>noscale <u>s</u>patiotemporal indexing <u>c</u>lustering, hereafter referred to as NASTIC, with DBSCAN and Voronoï tessellation clustering and demonstrated effective and efficient spatiotemporal resolution of molecular clusters.

## RESULTS AND DISCUSSION

### NASTIC workflow

We used a Python implementation of the R-tree spatial index https://pypi.org/project/Rtree/ incorporated into a framework utilizing the common Python modules SciPy, SciKit Learn, Numpy, Seaborn and Matplotlib. R-tree spatial indexing relies on use of rectangular bounding boxes rather than the irregular bounding regions encompassing a typical trajectory. For the purposes of spatiotemporal indexing, we therefore used an idealised rectangular bounding box based on the approximate radius of the convex hull (the boundary enclosing a series of points such that there are no concavities in the boundary line) encompassing the spatial extent of all the detections comprising a given molecular trajectory (Fig. 2A-D). The bounding box was extended into the time dimension allocating a user-defined time “thickness” that encompassed the duration of the tracked molecule (Fig. 2E). The resulting 3D (x,y,t) bounding box was indexed into a 3D R-tree database (Fig. 2F). To determine overlapping bounding boxes, each entry in the R-tree was queried, and returned a list of any other overlapping entries: e.g. entry A→[A,B], entry B→[B,A,C,D], entry C→[C,B,D], entry E→[E,F], entry F→[F,E] etc. Lists containing common entries were distilled down to lists representing clusters of overlapping bounding boxes: e.g. [A,B,C,D] and [E,F] (Fig. 2F), where each entry corresponds to a single molecular trajectory (Fig. 2G). A convex hull of the spatial extent of all detections associated with the clustered trajectories was used to define the cluster area. An important feature of NASTIC is that identification of temporally distinct clusters occupying the same spatial extent (Fig. 2H) is an intrinsic feature of the analysis. Querying the R-tree and defining spatiotemporal clusters is rapid, taking < 1 s for ∼5000 trajectories on a modest laptop.

**Figure 2.**
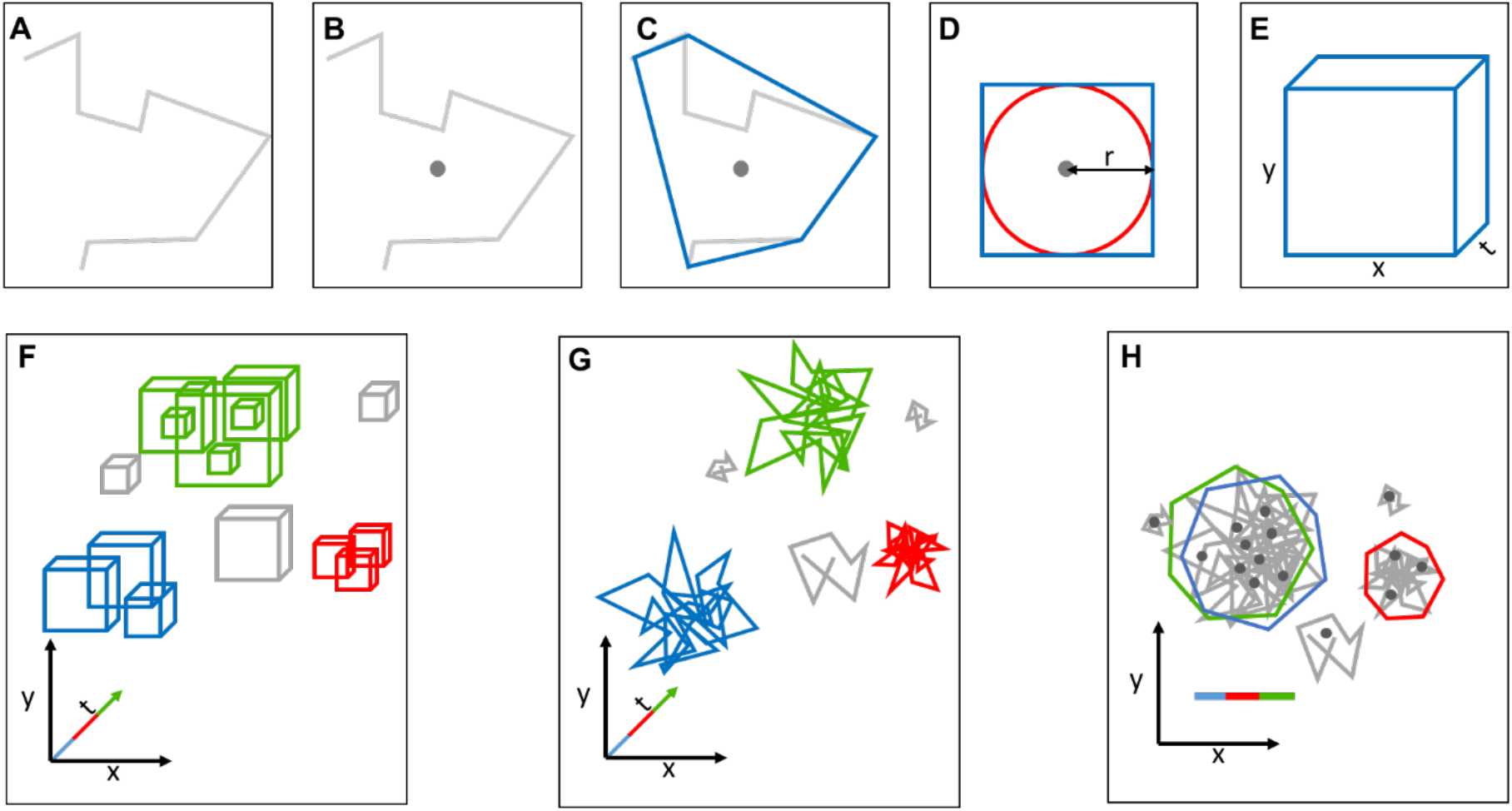
Schematic representation of NASTIC workflow. **(A)** Molecular trajectory composed of individual detections. (**B)** Spatiotemporal centroid representing the trajectory’s average position in space and time. (**C)** Convex hull (blue) defining the spatial extent of the trajectory. (**D)** Simplified 2D spatial bounding box (blue square) based on the approximate radius (r) of the convex hull (red circle). (**E)** 3D spatiotemporal bounding box of user-defined “thickness” in the time dimension. (**F)** R-tree spatiotemporal index of all trajectory bounding boxes. Discrete clusters of overlapping bounding boxes are indicated in color, and unclustered boxes in gray. (**G)** 3D clusters of trajectories associated with overlapping bounding boxes. (**H)** 2D representation of clustered trajectories. Colored polygons represent the spatial convex hull of all detections comprising each of the clustered trajectories. Clusters are colored according to the averaged detection time of their component trajectories, allowing assignment of overlapping clusters (green and blue) occupying the same spatial extent at different times.

### Optimum parameters for spatiotemporal clustering

The “holy grail” of a spatiotemporal clustering algorithm which works without operator input regardless of the density and distribution of SMLM data does not exist. Low density data decrease the likelihood of adjacent detections which can be considered clustered, while high density data conversely increase the likelihood of spurious clustering. Similarly, data in which detections are concentrated into multiple discrete areas within an acquisition window (e.g. axonal or dendritic neuronal projections in highly polarized cells) pose analytical challenges compared to data acquired across a single larger and flatter cell (e.g. a PC12 cell). Currently adopted spatial clustering algorithms require optimized user-defined parameters in order to best represent the molecular clustering dynamics of the data. For DBSCAN, the radius (ε) around each molecular centroid in which to scan for other centroids, and the minimum number of centroids within this radius (MinPts) to be considered a cluster, must be determined empirically. For Voronoï tessellation the threshold tile size below which a centroid is considered clustered must also be established against the average tile size or by using randomly distributed coordinates of similar density as the experimental data^15,16^. In all cases, the need to empirically determine optimized parameters for clustering exposes the potential for operator bias. In theory, NASTIC should assign trajectories into clusters based purely upon overlapping spatiotemporal boundaries, without parameters. In practice, two parameters are actually required: 1) a bounding box radius factor (*r*). The nature of R-tree indexing requires a rectangular bounding box for each trajectory. This simplified bounding box (Fig. 2D) only encompasses the full extent of the trajectory if its original convex hull (Fig. 2C) was perfectly circular. Applying a radius factor *r* > 1 is necessary to create a bounding box more representative of the full extent of the trajectory. 2) A time window (*t*), in seconds. This defines the temporal “thickness” of the bounding box (Fig 2E). Greater values of *t* increase the likelihood that spatially overlapping bounding boxes will also overlap in time to generate a discrete temporal cluster. A value of *t* equal to the 2x the acquisition time will in effect return purely spatial clustering, given that no trajectory can be considered temporally distinct from any other within this large acquisition time window.

We used simulated trajectory data with controlled density and spatiotemporal distribution for preliminary evaluation of a given clustering algorithm’s ability to designate clusters matching the ground truth inherent in the input data. We first generated a small dataset approximating the scale and density of a typical super-resolution acquisition at 50 Hz over 320 s. These data consisted of 50 randomly distributed trajectories within 4 μm^2^ and 0-320 s, interspersed with 20 discrete clusters, where each cluster contained 20 trajectories within a 0.1 μm radius and a 10 s time window. Each trajectory was a random walk with 8 - 30 segments of length < 0.1 μm with 20 ms between segments. A dataset was chosen which exhibited clusters of various types: (a) clusters discrete in space and time; (b) clusters which overlapped in space and time; (c) clusters which partially overlapped in space and time; and (d) clusters which overlapped in space but not in time. From the point of view of functional nanoclustering this last class of clusters are particularly important as they model functional hotspots on the plasma membrane which can repeatedly recruit molecules to a site of biological activity, such as the synaptic active zone ^21-23^.

Initially, NASTIC was performed using a limited range of *r* and *t* values, representative results of which are shown in Fig. S1. We found that *r* = 1.2 and *t* = 20 s returned spatial clusters most representative of the distribution of synthetic random walk trajectory data. Lower values of *r* and/or *t* returned multiple small spatiotemporal clusters for each of the discrete input clusters. Conversely, higher values returned fewer and larger clusters.

We expanded the exploration of the parameter space using a larger synthetic dataset consisting of 100 spatially distinct cluster regions randomly distributed on a simulated 100 μm^2^ membrane area. 20% of the regions constituted hotspots where 2-4 clusters occupied roughly the same spatial extent but occurred at different time points over the simulated 320 s acquisition. The final dataset thus consisted of 1095 trajectories in 140 spatiotemporally unique clusters with 7.82 ± 0.16 trajectories per cluster, with cluster radii of 74.86 ± 5.29 nm. The synthetic data also contained a background of 1000 randomly spatiotemporally distributed unclustered trajectories. NATSIC analyses were performed using a matrix of *r* (0.2 – 4.0) and *t* (1 – 640 s) values, and for each *r/t* pair the following metrics were evaluated: 1) number of trajectories in clusters, 2) the total number of spatiotemporally discrete clusters 3) the number of trajectories associated with each cluster and 4) the cluster radius. These metrics were compared against the ground truth values for the dataset and plotted as heatmaps of log2[observed:ground truth] for each *r/t* pair (Fig. S2). These plotted data reveal a complex relationship between *r, t* and each cluster metric, where lines of linked pale pixels highlighted the *r/t* pairs which returned metrics more closely matching the ground truth. We looked for the inflection point where these lines transitioned from vertical to horizontal. This generally occurred at *r* = 1.2 – 1.4 and *t* = 10 – 25 s, depending on the metric. When the pixels were averaged across the 4 metrics, the resultant plot (Fig. 3) demonstrated an inflection point around *r* = 1.2 - 1.4 and *t* = 15 – 20 s. In all subsequent NASTIC analyses, *r* = 1.2 and *t* = 20 s were therefore used as the default parameters.

**Figure 3.**
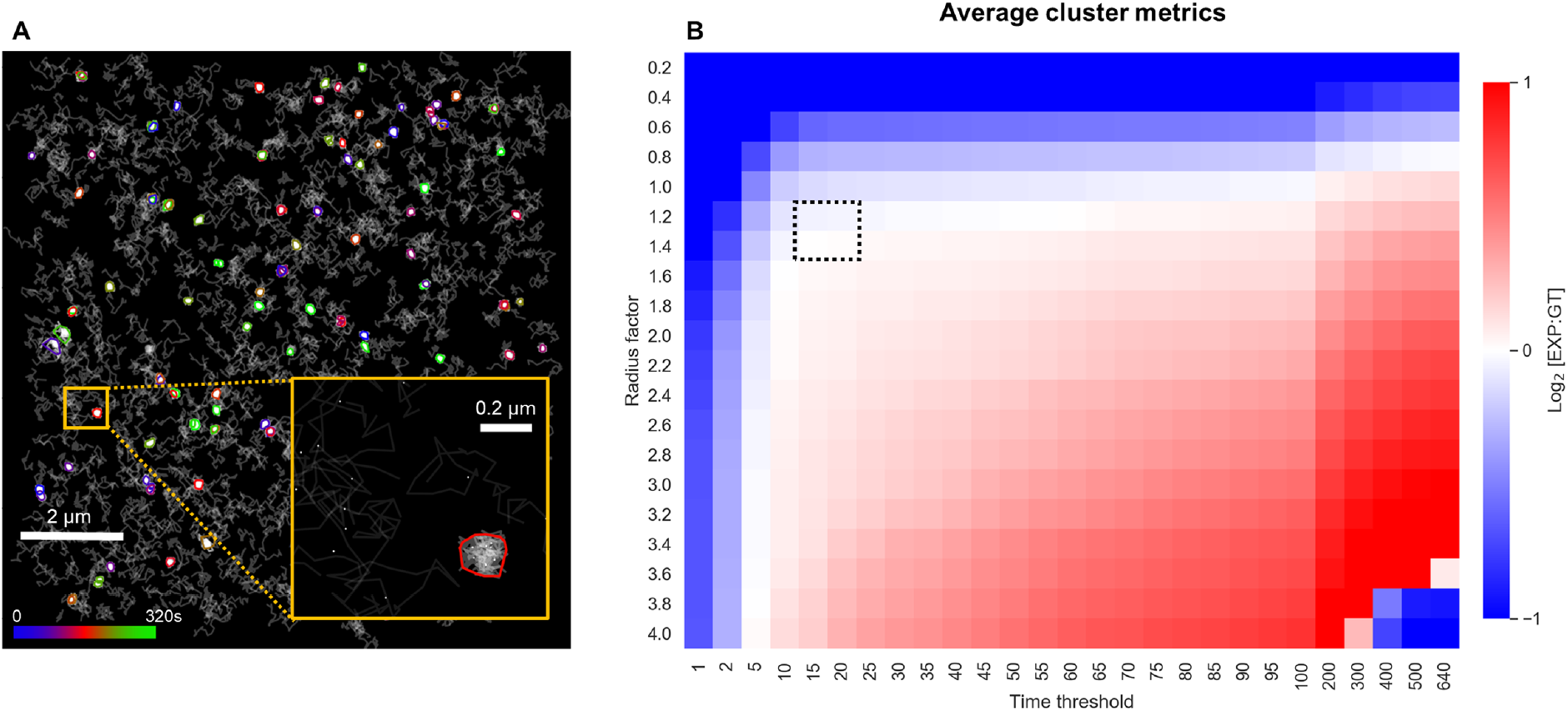
Effect of bounding box radius and time window on spatiotemporal clustering metrics. *In silico* random walk trajectory data consisting of 1095 trajectories in 140 spatiotemporally unique clusters with 7.82 ± 0.16 trajectories per cluster, with cluster radii of 74.86 ± 5.29 nm. The data also contained a background of 1000 randomly spatiotemporally distributed unclustered trajectories. Clusters are randomly distributed within a simulated 320 s acquisition window. **(A)** Nanoscale spatiotemporal indexing (NASTIC) of this simulated trajectory data using *r* = 1.2, *t* = 20 s. Cluster boundaries represent the extent of the detections associated with clustered trajectories, and are colored according to the average detection time. The inset displays a zoomed view of a single cluster against a background of unclustered trajectories, with trajectory centroids indicated with a dot. **(B)** Heatmap of averaged metrics (cluster number, cluster radius, trajectories per cluster and number of clustered trajectories, see Fig. S2). Each pixel represents the average log_2_ ratio of the experimental observed (EXP) value for a given *r/t* pair to the ground truth (GT). Ratios < -1 and > 1 are displayed as 1 and -1 respectively. Pale regions indicate *r/t* pairs which return cluster metrics close to the ground truth. The approximate inflection point where the pale line transitions to horizontal is indicated with a dotted box.

### Comparison of clustering algorithms using simulated trajectory data

Having established that NASTIC could resolve clusters in simulated data, we sought to compare spatiotemporal indexing with density-based and segmentation-based clustering on the same data. Using the smaller synthetic dataset described above, we performed a qualitative comparison of NASTIC using optimized parameters (*r* = 1.2, *t* = 20 s) against two widely used spatial clustering algorithms, DBSCAN and Voronoï tessellation. For fair comparison purposes the clustering comparison was achieved via similar Python frameworks differing only by the commonly used Python modules implemented for clustering: SciKit Learn DBSCAN and SciPy.Spatial Voronoi. For DBSCAN, the centroids of all trajectories were analysed using ε **=** 0.055 μm and MinPts = 3, which were chosen to return spatially distinct clusters of similar dimensions to the input data (radius ∼0.1 μm). Voronoï tessellation of all trajectory centroids was thresholded such that tiles with an area of < 0.004 μm^2^ were considered clustered. This value was empirically determined to best yield clusters reflective of the input data. Clustered tiles were grouped into discrete clusters if they shared one or more edges with other clustered tiles. In all three cases (NASTIC, DBSCAN and Voronoï), a cluster was defined as three or more proximal centroids. The spatial extent of each cluster was determined by a convex hull of all the trajectory detections associated with the cluster, and the trajectories, centroids and clusters visualized by Python Matplotlib (Fig. 4).

**Figure 4.**
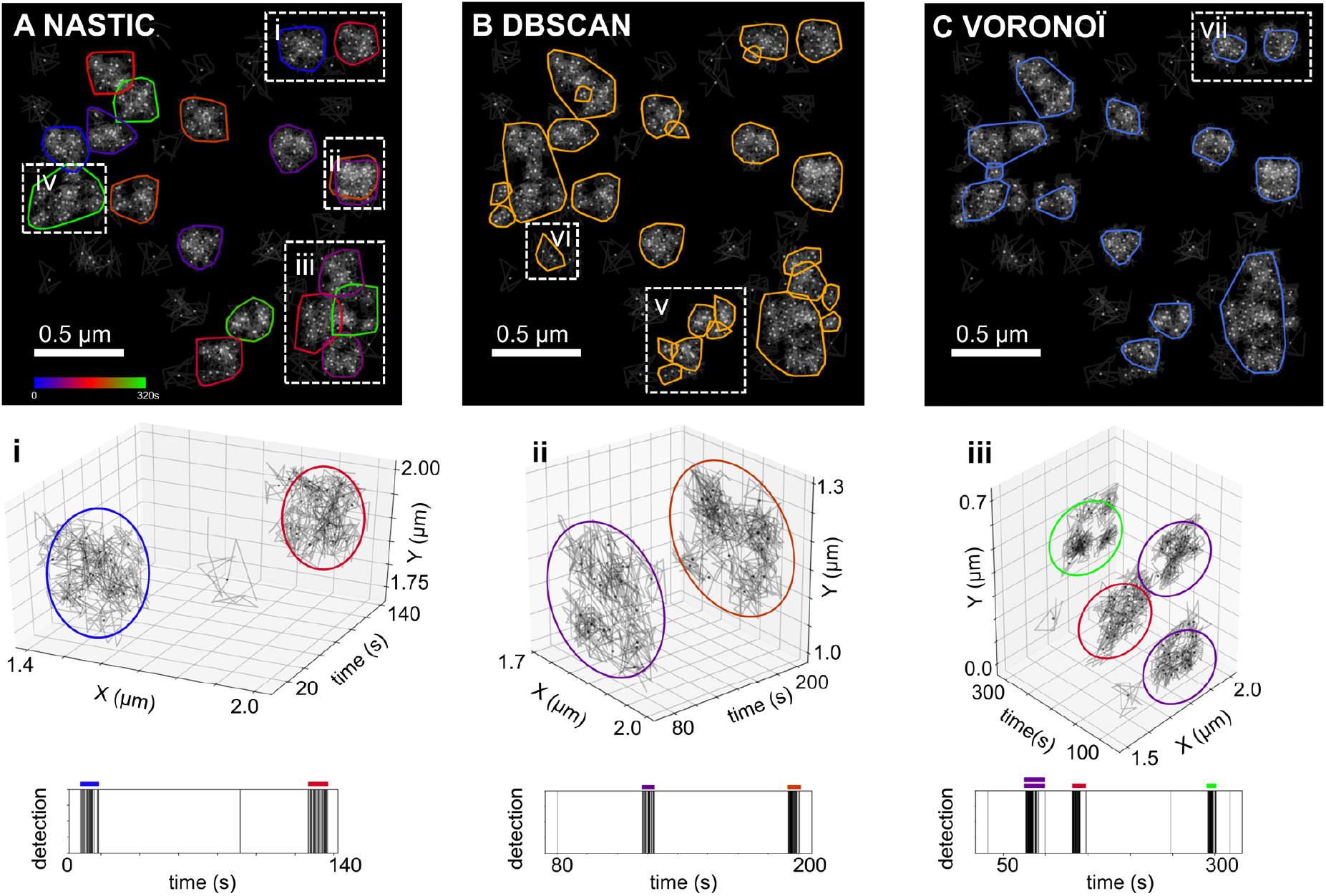
Resolution of spatiotemporal clusters in simulated data. *In silico* random walk trajectory data consisting of 20 equivalently sized spatiotemporal clusters with 20 trajectories per cluster and a background of 50 non-clustered trajectories. Clusters are randomly distributed within a simulated 320 s acquisition window. The spatial centroid of each trajectory is represented as a dot. (**A)** Clustering using NASTIC using *r* = 1.2, *t* = 20 s. Trajectories are considered as clustered if their bounding boxes overlap with other trajectory bounding boxes within a 20 s time window. Cluster boundaries represent the extent of the detections associated with clustered trajectories, and are colored according to the average detection time. Insets highlight different classes of clustering: (**i)** distinct clusters resolved in space and time; (**ii)** spatially overlapping clusters resolved in time; (**iii)** clusters with a degree of spatial and temporal overlap; (**iv)** clusters which overlap in space and time. 3D (x, y, t) projections of highlighted clusters i – iii and the associated detection times (lower panels) demonstrate distinct temporal clustering. (**B)** DBSCAN spatial clustering using **ε =** 0.055 μm and MinPts = 3. (**C)** Voronoï tessellation spatial clustering. Trajectories with an associated Voronoï tile area < 0.004 μm^2^ were considered clustered. In all analyses a cluster is defined as 3 or more proximal centroids.

NASTIC was able to resolve the 20 clusters input data into 18 distinct clusters of approximately equal size consistent with the 0.1 μm random distribution radius used to generate each cluster (Fig. 4A). Clusters were observed which represented: (Fig. 4A i) distinct clusters resolved in space and time; (Fig. 3A ii) spatially overlapping clusters resolved in time; and (Fig. 4A iii) clusters with a degree of spatial and temporal overlap. A single larger cluster (Fig. 4A iv) was also observed which was consistent with two clusters overlapping in both time and space, accounting for the remaining 2 clusters in the input data. 3D (x, y, t) projections of the data demonstrated that spatiotemporal indexing resolved the clusters that were distinct in both space and time. Trajectories which occupied the same spatial extent (Fig. 4A ii) could be resolved into discrete spatiotemporal clusters, as could trajectories with some degree of spatiotemporal overlap (Fig. 4A iii). Importantly, DBSCAN and Voronoï tesselation, neither of which can natively resolve in the temporal dimension, failed to separate these overlapping clusters, reporting overlapping and proximal (touching) spatiotemporal clusters as single spatial clusters. Further, in some cases DBSCAN resolved areas of higher density within a spatial cluster into smaller clusters (Fig. 4B v). DBSCAN also reported areas of background “noise” as clusters (Fig. 4B vi). When compared to both NASTIC and DBSCAN, Voronoï tessellation returned slightly smaller clusters as a result of centroids on the edge of a cluster with Voronoï tiles larger than the 0.004 μm^2^ threshold (Fig. 4C vii).

We expanded the comparison by creating 10 randomised synthetic datasets based upon the same seed conditions as described for our exploration of *r/t* values on clustering metrics. These datasets consisted of 100 spatially distinct cluster regions randomly distributed on a simulated 100 μm^2^ membrane area. 20% of the regions constituted hotspots where 2-4 clusters occupied roughly the same spatial extent but occurred at different time points over the simulated 320 second acquisition. Each dataset thus consisted of approximately 140 spatiotemporally unique clusters of 6-10 trajectories per cluster, with cluster radii of approximately 75 nm. Clusters constituted approximately 1100 trajectories, against a background of 1000 randomly spatiotemporally distributed unclustered trajectories. The 10 datasets were analysed using NASTIC (*r* = 1.2, *t* = 20 s), DBSCAN (**ε =** 0.05 μm, MinPts = 3) and Voronoï tessellation (tile threshold 0.1 μm^2^). For all algorithms the parameters were determined empirically to optimise returned metrics corresponding to the ground truth of the input synthetic data.

As shown in Fig. 5, NASTIC consistently returned metrics most closely matching the ground truth of the simulated data. Although DBSCAN and NASTIC both reported similar numbers of trajectories within clusters, DBSCAN was unable to resolve the clusters in the hotspots and therefore reported cluster numbers closely matching the 100 input cluster regions. This also resulted in DBSCAN reporting higher average numbers of trajectories in a cluster, and a larger average cluster size due to spatially overlapping but temporally distinct clusters being treated as single larger spatial clusters. Voronoï tessellation consistently reported fewer clustered trajectories, and fewer and smaller clusters, with fewer trajectories within each cluster. In our hands, both NASTIC and DBSCAN can be considered to return data reflecting the spatial clustering of the trajectories, with NASTIC natively returning more accurate data reflecting the unique spatiotemporal clustering of the trajectories. The comparatively poor results obtained by Voronoï tessellation clustering may reflect issues of the algorithm to accurately segment lower density trajectory centroid information, as opposed to higher density detection information. In any case, Voronoï tessellation could not distinguish temporally distinct clusters and would be expected at best to match the metrics returned by DBSCAN.

**Figure 5.**
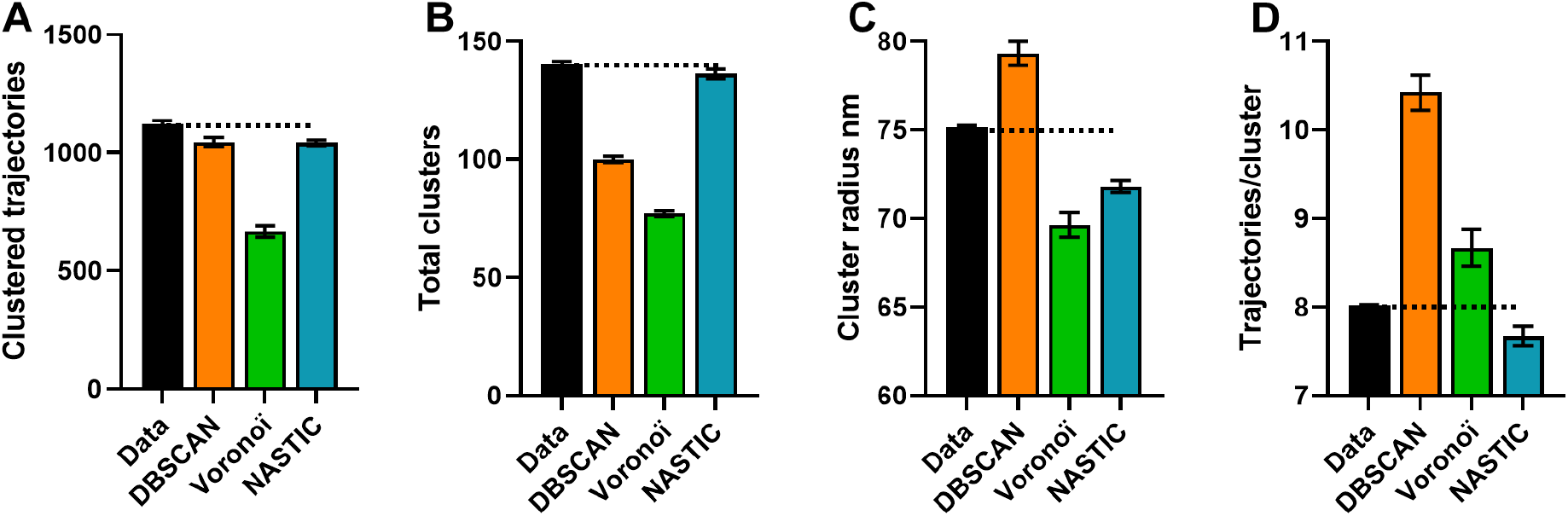
Comparison of clustering algorithms using synthetic data. Data simulating multiple 50Hz acquisitions over 320 s as described in the text. (**A)** Total trajectories in clusters, (**B)** Total unique clusters, (**C)** Average cluster radius (nm) and (**D)** Average trajectories in a cluster. Black bars represent the ground truth in the simulated data, colored bars represent the metrics returned by DBSCAN (ε = 0.05 μm, MinPts = 3, orange), Voronoï tessellation (tile threshold 0.01 μm^2^, green) and NASTIC (*r* = 1.2, *t* = 20 s, blue). Error bars show standard error of the mean (SEM) across 10 datasets. The dotted black line shows the average value in the inputted synthetic data.

### Clustering analysis of syntaxin1a-mEos2 super-resolution imaging data

While simulated data offer the ability to precisely model trajectory density and clustering, the molecular environment within a living cell is far more varied and dynamic and represents a greater analytical challenge. We therefore next applied NASTIC, DBSCAN and Voronoï tessellation clustering to single particle tracking photoactivated light microscopy (sptPALM^1,24-26^) data obtained from syntaxin1a (Sx1a) tagged with mEos2 in live neurosecretory PC12 cells (Fig. 5A - C). Sx1a is a member of the SNARE protein family that is located on the plasma membrane of neurons and neurosecretory cells and is involved in mediating synaptic and neurosecretory vesicle fusion^6,11,27-29^. As anticipated, all three techniques were able to detect spatial clustering of Sx1a, with NASTIC also able to natively assign temporal information to the clusters.

In the 100 μm^2^ region of interest analysed, 2564 trajectories were selected, of which 1116 were assigned based on spatiotemporal indexing to 172 clusters. The average number of trajectories associated with each cluster was 6.592 ± 0.347 and the average radius of each cluster was 84.587 ± 2.329 nm. On average a cluster existed for 12.15 ± 0.87 s. In addition to discrete spatiotemporal clusters (Fig. 6D i), we identified a total of 27 overlapping clusters (conservatively defined as those whose centroids are separated by less than 0.5 of the average cluster radius) (Fig. 6D ii) where Sx1a molecules appeared to be repeatedly recruited to the same region of the plasma membrane. Plotting the trajectories of the overlapping and discrete clusters allowed visualization of 17 potential functional hotspots for Sx1a clustering on the PC12 cell plasma membrane ^6,12,30-33^ (Fig. 7A, B). As evidenced by mean square displacement (MSD) curves for unclustered and clustered trajectories (Fig. 7C), these clusters truly represent nanomolecular assemblies where Sx1a was constrained into lower mobility states. If these clusters were merely artifacts of randomly overlapping trajectories, the mobility of clustered and unclustered trajectories would be expected to be similar.

**Figure 6.**
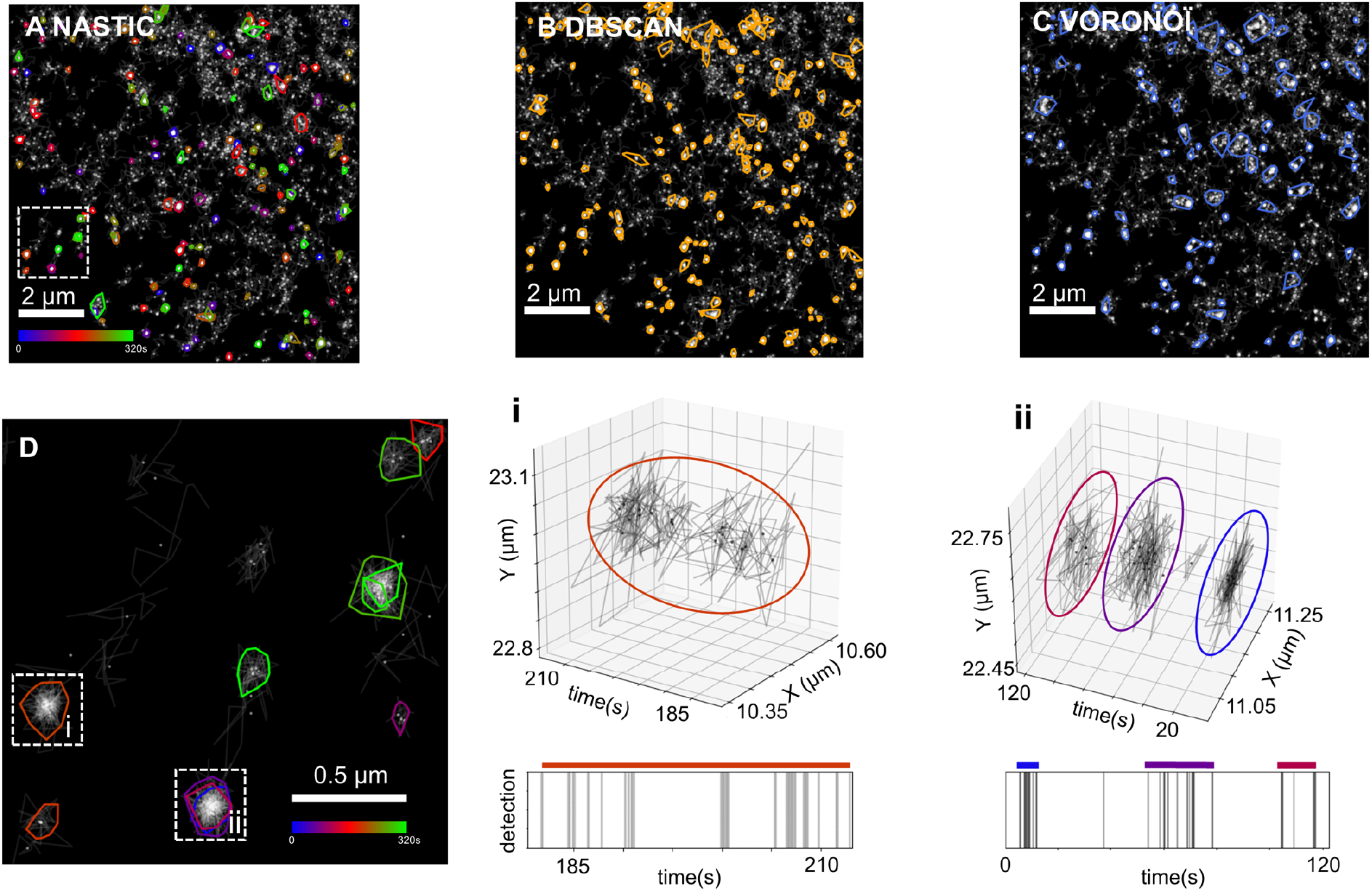
Resolution of spatiotemporal clustering in live-cell molecular trajectory data. Sx1a-mEos2 sptPALM data acquired at 50 Hz over 320 s. Clustering based on (**A)** NASTIC using ***r*** = 1.2, ***t*** = 20 s, (**B)** DBSCAN using **ε =** 0.055 μm and MinPts = 3 and (**C)** Voronoï tessellation using tile threshold 0.015 μm^2^. (**D)** Magnification of the region indicated by the dotted box in (A). Cluster boundaries represent the extent of the detections associated with clustered trajectories, and are colored according to the average detection time. Insets highlight different classes of clustering further visualized by 3D (x, y, t) projections of highlighted clusters and the associated detection times: (**i)** single spatiotemporal cluster; (**ii)** spatially overlapping clusters resolved in time.

**Figure 7.**
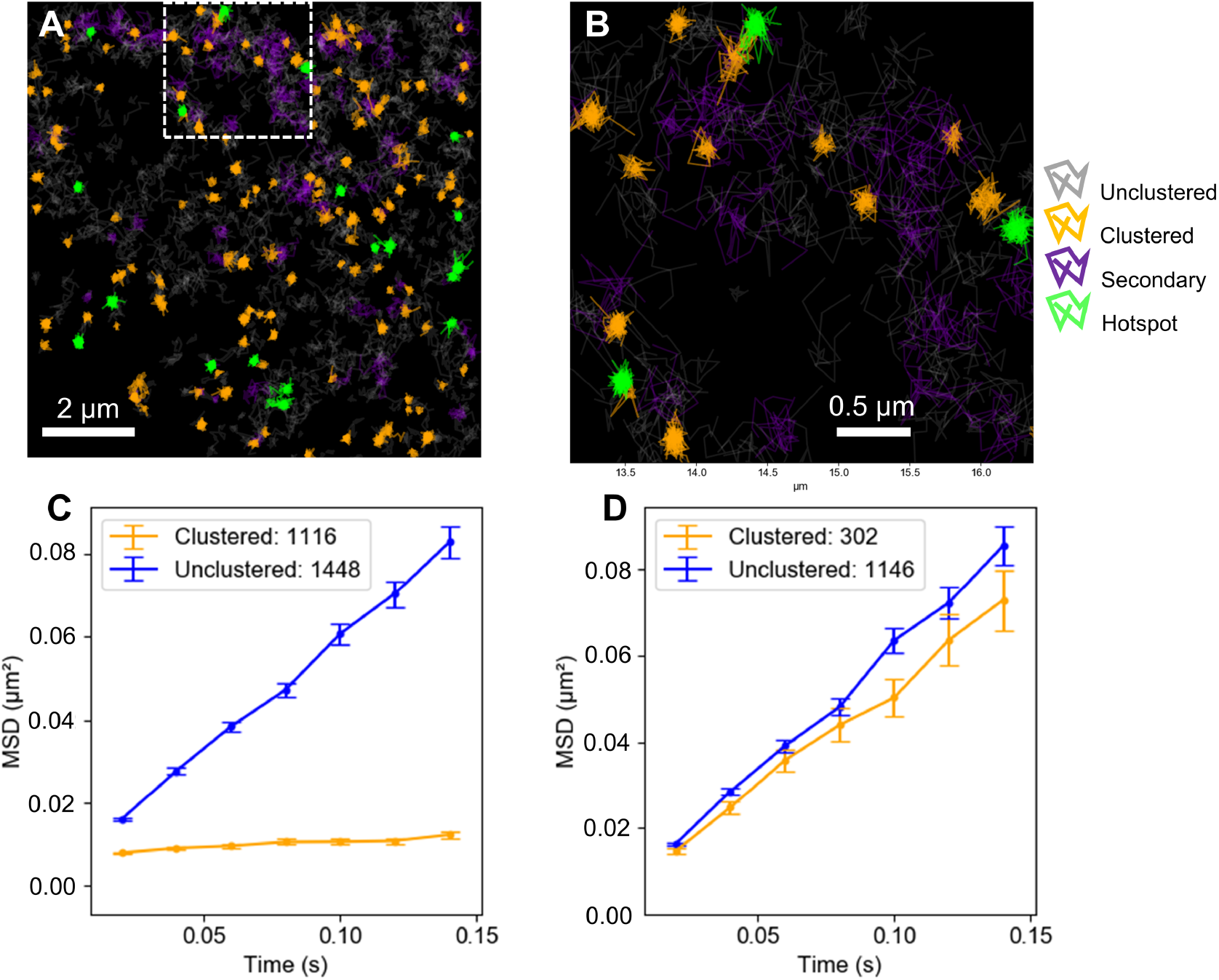
Identification of time resolved (primary) and non-time resolved (secondary) clusters reveals clustering hotspots. Sx1a-mEos2 sptPALM data acquired at 50 Hz over 320 s. (**A)** Analysis using spatiotemporal indexing using ***r*** = 1.2, ***t*** = 20s. Unclustered trajectories are shown in grey, spatially discrete clustered trajectories are shown in orange, trajectories belonging to spatially overlapping clusters are shown in green. Secondary analysis of the unclustered trajectories from the primary analysis, using ***r*** = 1.2, ***t*** = 640. Trajectories in secondary clusters are shown in purple. Dotted box indicates the area enlarged in (**B)**. (**C, D)** Mean square displacement (MSD) curves of clustered and unclustered detections from the primary analysis and secondary analysis respectively. Each point represents the average MSD of the indicated number of trajectories. Error bars indicate the standard error of the mean (SEM).

### Identification of loosely interacting trajectories by iterative clustering

The 172 “primary” time resolved Sx1a clusters identified by NASTIC above represented those whose individual trajectories overlapped within a 20 second time window. We next sought to identify and characterize any remaining “secondary” clusters representing trajectories which overlapped at any time within the 320 s acquisition window. The 1448 unclustered trajectories from the primary analysis described above were therefore reanalysed using *r* = 1.2 and *t* = 640 s (Fig. 7A, B). A further 302 trajectories were assigned to 66 individual secondary clusters. Rather than discrete circular nanomolecular structures, these secondary clusters tended to represent unconfined trajectories whose larger bounding boxes increased the likelihood of overlap during the course of the acquisition. This is evidenced by the MSD curve for these secondary clusters (Fig. 7D) which show dramatically less mobility difference between unclustered and secondary clustered trajectories. The generally diffuse nature of the secondary clustered trajectories strongly contrasts with the dense compact circular nature of the time-resolved primary clusters (Fig. 7B) and lends further support to the ability of time resolved spatiotemporal indexing to identify true nanomolecular clustering against a significant background of higher mobility unclustered trajectories.

### Characterization of spatiotemporal hotspots

NASTIC allows native identification of hotspots where spatial clusters form, dissociate, and reform more than once over the acquisition period. With sufficient clusters identified, additional metrics related to spatiotemporal clustering may be defined. These metrics were obtained using DBSCAN on the spatial centroids of the 172 clusters identified above, in essence treating hotspots as clusters of clusters. DBSCAN was performed using MinPts = 2 and ε values in the range 0 – average cluster radius. **Overlap probability** (Fig. 8A) measures the likelihood of a cluster centroid having another cluster centroid within a given distance. This is computed as `1 – (unclustered centroids/total centroids)`. At very small values of ε, the chance of any cluster having a proximal cluster is low, as these clusters must occupy essentially the same spatial extent. Conversely, at a distance corresponding to the average cluster radius, the likelihood of a proximal cluster increases, as these clusters essentially need only to touch edges. To determine the degree to which cluster overlap was driven by biological distribution rather than chance, we performed a Monte Carlo simulation (N=50), using 172 clusters randomly distributed over the same 100 μm^2^ analysis area (Fig 8A, red plot). The result of this analysis demonstrates that random cluster overlap contributed little to the observed overlap. **Hotspot membership** (Fig. 8B) defines the average number of clusters detected in each hotspot. **Intercluster time** (Fig. 8C) measures the average time between clusters in each hotspot. Together these analyses show that there was an approximately 17% chance of any given Sx1A cluster forming a hotspot with another cluster within 41.5 nm (half the average cluster radius). The average hotspot contained between 2-3 clusters, and the average time between each cluster in a hotspot was 45 s. **Cluster number** (Fig. 8D), measured as the number of discrete spatiotemporal clusters per μm^2^, varied over the duration of the acquisition but did not dramatically trend up or down. This is indicative of a potential “steady-state” of clustered Sx1a on the plasma membrane. As shown in Figs 8E-H, NASTIC was clearly able to identify regions on the plasma membrane (highlighted boxes) where new trajectories were recruited into clusters by lateral trapping over many tens of seconds. Through the use of a user-defined time window (*t*), NASTIC could assign these multiple trajectories into discrete temporal clusters which then allowed the additional hotspotting metrics described above. The functional significance of these hotspots is currently unknown but could represent docking/priming sites at the plasma membrane^34^.

**Figure 8.**
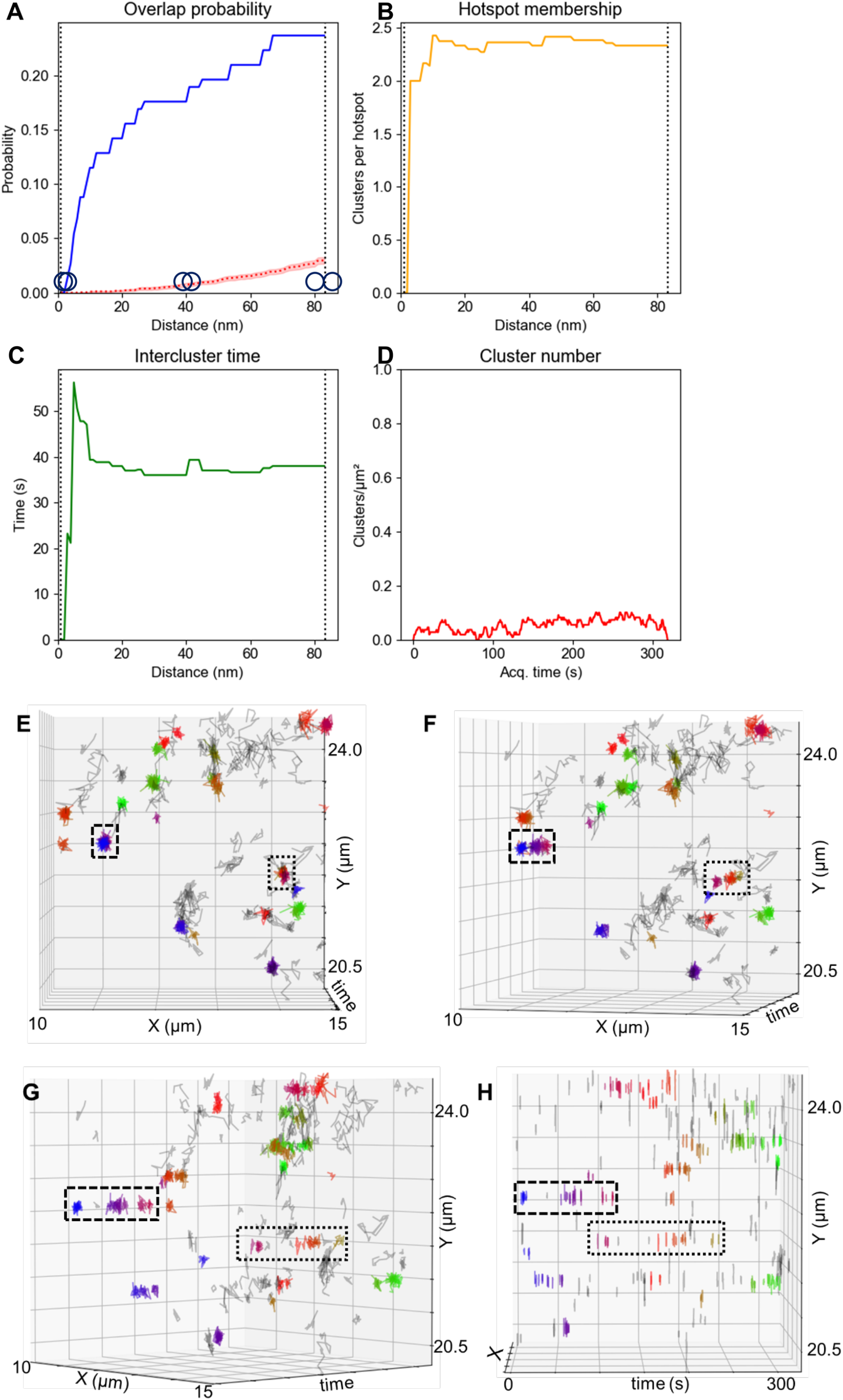
NASTIC spatiotemporal metrics. Sx1a-mEos2 sptPALM data acquired at 50Hz over 320 s analysed by NASTIC. (**A)** Probability of cluster overlap using DBSCAN of cluster centroids identified by NASTIC, **ε =** 0.001 - 0.083μm (average cluster radius) and MinPts = 2. Monte Carlo simulation (N=50) using 172 randomly distributed cluster centroids was used to establish the degree of random overlap of clusters of the same number and density as the experimental data. The dotted red line indicates the average overlap probability, translucent red indicates the standard error of the mean. The left and right dotted vertical lines represent 0.001 μm and 0.083 μm respectively. At 0.001 μm two clusters must essentially completely overlap to be considered as a hotspot, as illustrated by the overlapping circles. At 0.083 μm, two clusters are considered as members of a hotspot if their edges touch, as indicated pictorially by the two touching circles. (**B)** Average number of clusters in a hotspot, as a function of distance. (**C)** Average time between clusters in a hotspot, as a function of distance. (**D)** Number of unique spatiotemporal clusters observed at 1 second intervals over the 320 s acquisition. (**E-H)** Progressive rotation around the y-axis of the 3D [x,y,t] projection of spatiotemporal Sx1a-mEos2 trajectory data, with clustered trajectories indicated in color. Dotted and dashed boxes show representative single spatial clusters which resolve into multiple spatiotemporal clusters (hotspots). The dashed boxes correspond to the spatiotemporal hotspot observed in Fig. 6d (ii).

### Using spatiotemporal indexing to define activity-dependent changes in nanoclustering dynamics

A growing body of literature has demonstrated that the spatial and temporal nanoscale organization of key proteins in the neuroexocytic pathway can change in an activity-dependent manner. These changes may reflect the functional clustering of a range of proteins required for the synaptic vesicle docking, priming, fusion and recycling at the heart of neuronal synaptic communication^35^. They may also reflect activity-dependent protein conformational changes which alter the protein’s homo- / heterodimerization^36^. Analysis of molecular trajectory data obtained from SMLM experiments can provide insights into aggregate changes in the mobility of molecules, using metrics such as MSD as an indirect measure of the degree of their potential confinement in nanomolecular clusters. These can be further expanded using statistical techniques such as Hidden Markov Modelling (HMM^37,38^) to partition a molecular population into mobility states and transitions between them^9,39,40^, and Ripley’s K functions^10,17,41^ to derive insights into the point dispersion and cluster size. More directly, clustering analysis can reveal pertinent metrics related to the number and size of clusters, the number of molecules within clusters, and their lifetimes and rates of formation. These metrics can be averaged across sufficient datasets and compared between experimental conditions, such as non-stimulated and stimulated cells, to assign statistical significance to the degree of change. As demonstrated herein, NASTIC allows significant expansion of nanocluster analysis to include metrics such as the extent of molecular clustering hotspots, the number of temporally distinct clusters within these hotspots and the time between these clusters. We sought to establish the degree to which NASTIC could generate statistically significant measures of the change in nanocluster dynamics of another key priming protein, Munc18-1 tagged with mEos2, in response to stimulated exocytosis in PC12 cells. Accordingly, sptPALM data was acquired from 9 cells under unstimulated conditions, and following stimulation with BaCl2 (2 mM). NASTIC analysis was performed on individual cells. For each cell, the metrics for each cluster were generated, together with average metrics for all clusters in the cell. Two comparison analyses were then performed: (1) the metrics for the 2610 pooled clusters detected across 9 unstimulated cells were compared with those from 2588 pooled clusters observed across the 9 stimulated cells (Fig. 9), and (2) the averaged metrics from 9 unstimulated cells were compared with those from 9 stimulated cells (Fig. S3). These analyses suggested that secretagogue stimulation of PC12 cells resulted in significantly smaller clusters of Munc18-1. This is in agreement with our previously published work using autocorrelation of fast Fourier transformed image data^11^ demonstrating that Munc18-1 exits the confinement of nanocluster in response to stimulation following opening and engagement of cognate Sx1A in SNARE complex formation. The rate of detection of new trajectories over the lifetime of the cluster appeared higher in the stimulated cells, which might reflect activity-dependent changes to the dynamics of molecular recruitment into clusters.

**Figure 9.**
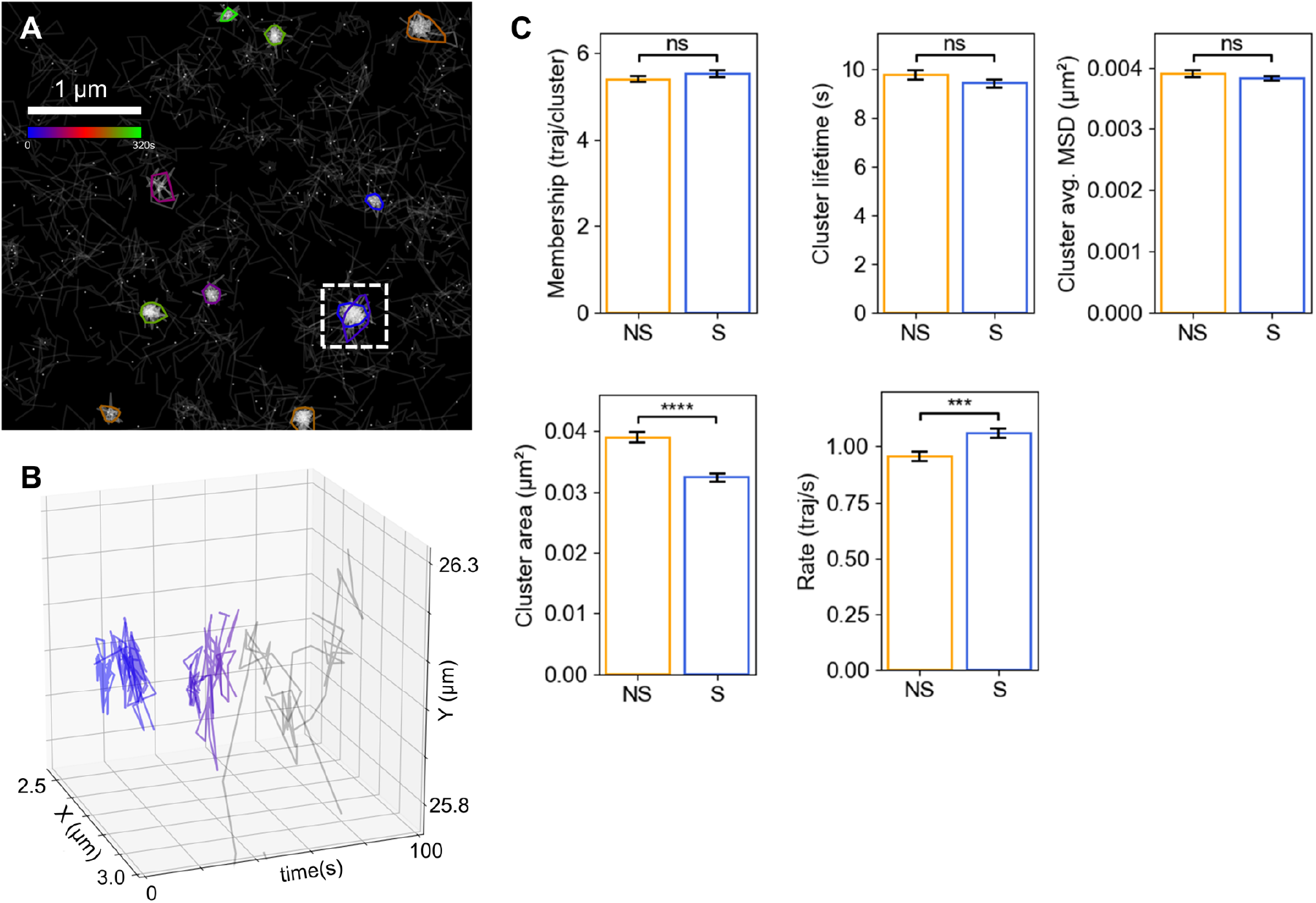
Difference in cluster metrics in response to stimulated exocytosis. Munc18-1-mEos2 sptPALM data acquired from unstimulated PC12 cells (N=9) and PC12 cells stimulated with 2 mM BaCl_2 (_N=9). (**A)** Representative image of clusters identified using spatiotemporal indexing with ***r*** = 1.2, ***t*** = 20 s. Dotted box highlights an overlapping cluster hotspot (**B)** 3D (x, y, t) projection of highlighted hotspot clusters. (**C)** Comparison of indicated cluster metrics of 2610 pooled clusters from unstimulated cells and 2588 clusters from stimulated cells. The significance of the difference between conditions was determined by unpaired *t*-test (ns = no significance, *** = p < 0.001, **** = p < 0.0001). NS = no stimulation, S = stimulation with 2 mM BaCl_2_.

### NASTIC of individual trajectory segments vs whole trajectory bounding boxes

Spatiotemporal indexing of trajectory bounding boxes allows the determination of clusters of overlapping trajectories which potentially interact in space and time. However, the more precise locations within the cluster where the overlap occurs are not recovered. As schematically represented in Fig. 10A, the boundary of a NASTIC cluster represents the furthest extent of the individual detections of all the overlapping trajectories in the cluster. Within this cluster, individual segments of each trajectory (a segment being defined as the line connecting a molecular detection with its subsequent detection) will overlap with segments from other trajectories. These represent regions within the cluster where there is an increased likelihood of the molecular overlap. A threshold can be determined based on the degree of segment overlap, beyond which the segments themselves can be considered as clustered. In scenarios of very high trajectory density and/or very long trajectories, determining clusters solely on overlapping trajectories (or DBSCAN/Voronoï tessellation of trajectory centroids) may lead to large clusters of low segment density (as demonstrated in Fig. 7B) which do not truly reflect potential molecular overlaps. We therefore investigated the application of spatiotemporal indexing to the bounding boxes of each individual trajectory segment (segNASTIC) in an effort to gain more fine-grained clustering information in high trajectory density data. Each trajectory segment was assigned a bounding box comprising its x and y extent, with a time “thickness” as described above. Trajectory segment bounding boxes were indexed into a 3D R-tree, which was queried to generate lists of potentially spatiotemporally overlapping segments. From these lists, the degree of overlap of each segment with segments from other trajectories was determined. Across all segments in an acquisition, a histogram was generated showing that the majority of trajectory segments had low overlap (Fig. S4). The inflection point of the histogram (the average segment overlap) was chosen as automatic threshold beyond which a segment was considered as potentially clustered. Clusters of thresholded overlapping segments were derived, representing more tightly defined areas on the plasma membrane where molecules were confined to interacting lower mobility states. We examined the benefit of this approach using data acquired by uPAINT (universal point accumulation for imaging in nanoscale topography ^42^) analysis of PC12 cells expressing Sx1a-EGFP (C-term tag) tracked by Atto-647-labelled anti-GFP nanobodies applied extracellularly ^11^. uPAINT acquisitions generally result in a higher density of relatively longer and more diffuse trajectories when compared to sptPALM, as they use organic dyes which are brighter and less prone to photobleaching. As demonstrated in Fig. 10B, spatiotemporal indexing of whole trajectory bounding boxes successfully identified clusters of confined trajectories in the uPAINT data, as well as a number of clusters consisting of relatively diffuse trajectories. The spatiotemporal indexing of trajectory segment data was represented as a pseudo-density map of trajectory overlap (Fig. 10C), which clearly showed regions of higher potential molecular interaction. The convex hull of the detections in each group of overlapping thresholded segments was used to define a unique spatiotemporal cluster. The trajectories associated with the clustered segments were colored, which enabled us to demonstrate the disparity between the size of the segment clusters and the extent of their parent trajectories (Fig. 10D). Compared with trajectory clustering, segment clustering generally returned smaller more tightly defined clusters, and far fewer clusters of trajectories with diffuse segments.

**Figure 10.**
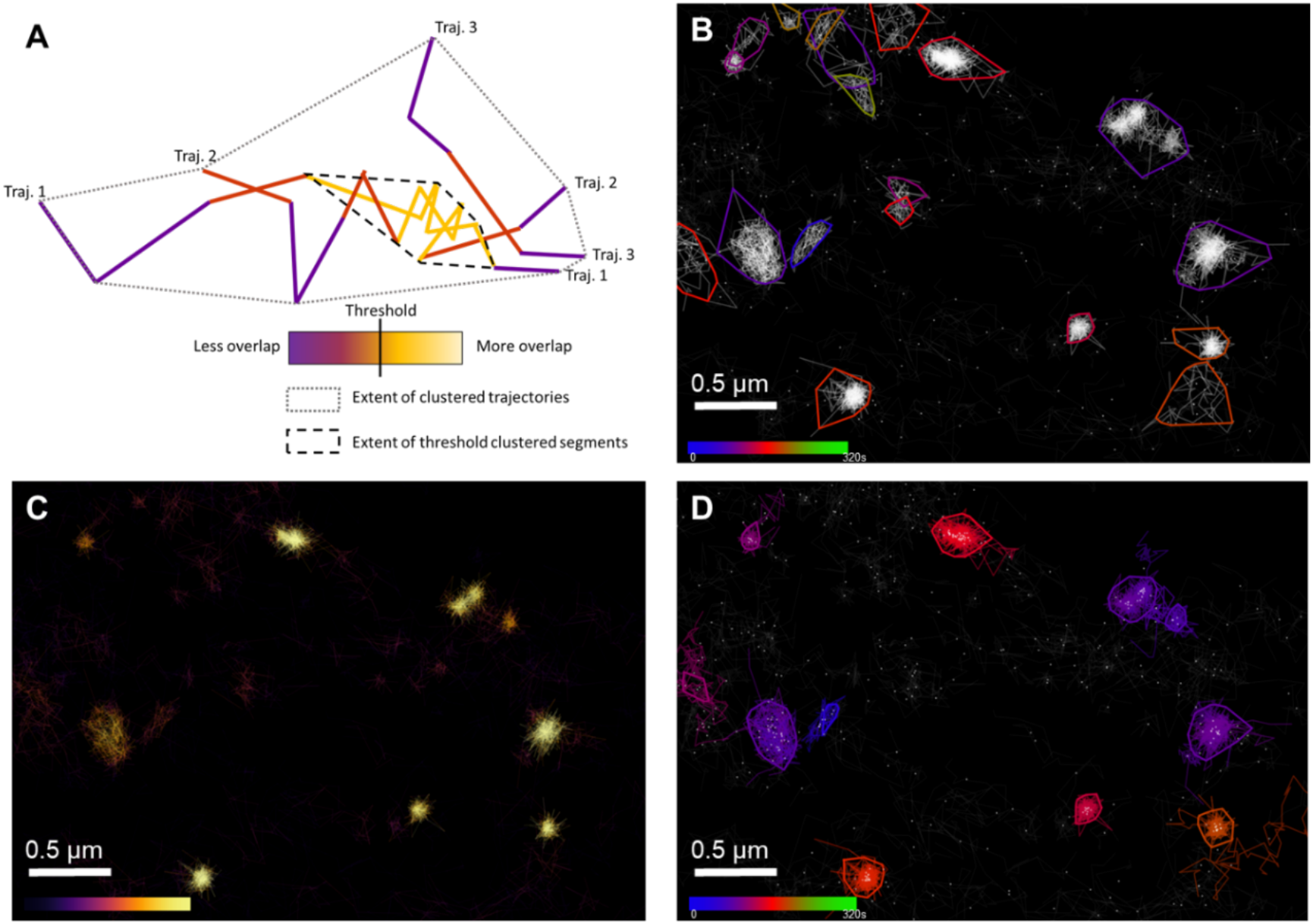
NASTIC of trajectory segments (segNASTIC). **(A)** Schematic representation of trajectory segment thresholding, based on overlap with segments from other trajectories. (**B)** Sx1a-EGFP imaged by uPAINT using Atto-647-labelled anti-GFP nanobodies in PC12 cells. Spatiotemporal clusters identified using spatiotemporal indexing of trajectory bounding boxes using ***r*** = 1.2 and ***t*** = 20 s. Each colored cluster boundary represents the convex hull of the detections belonging to all trajectories in the cluster (**C)** Pseudo-density map of trajectory segment overlap, with each trajectory colored according to the number of overlaps with other trajectory segments, as determined by spatiotemporal indexing of segment bounding boxes. (**D)** Spatiotemporal clusters identified using thresholded segments ***t*** = 20 s. Each colored cluster represents the convex hull of detections belonging to the clustered segments. All trajectories containing clustered segments are shown in the same color as the cluster.

However, this additional resolution does come at the price of the increased computational overhead of creating and querying a spatial index with 10-100 as many bounding boxes, resulting in an approximate doubling of the total analysis time when compared to NASTIC (Table 1). To compare the relative analysis times of NASTIC, segNASTIC, DBSCAN and Voronoï tesselation we used these pipelines to analyse a dataset consisting of 4207 trajectories (a subset of the Sx1a-mEos2 data presented in Figs. 6 and 7).

**Table 1.** Relative clustering analysis times. For each of the clustering pipelines, the time (s) to complete each stage of the analysis of the same dataset containing 4207 molecular trajectories is indicated. The analysis time does not include the time taken to visualise and display the clustered data.

| ANALYSIS TIME | NASTIC | segNASTIC | DBSCAN | VORONOÏ |
| --- | --- | --- | --- | --- |
| Selecting detections in ROI | 1.05 | 1.05 | 1.17 | 1.11 |
| Trajectory metrics (including bounding boxes) | 6.29 | 3.697 | 2.74 | 2.45 |
| Clustering | 0.49 | 13.34 | 0.02 | 12.7 |
| Cluster metrics | 1.93 | 1.402 | 1.72 | 1.36 |
| TOTAL | 9.76 | 19.489 | 5.65 | 17.62 |

In our hands, DBSCAN was capable of performing the clustering of trajectory detections faster than any other methods. However, NASTIC natively delivered additional temporal clustering information that neither DBSCAN nor Voronoï tessellation could provide. Interestingly, segNASTIC, which in addition to temporal information also provides increased spatial cluster resolution in very high-density data, has an analysis time comparable to purely spatial clustering by Voronoï tesselation.

Having demonstrated that segNASTIC returns smaller clusters representing the truly overlapping regions of each trajectory, we next compared the metrics returned from analysis of the complete Sx1a-mEos2 dataset (17598 trajectories) by NASTIC and segNASTIC as examined in Table 1. As shown in Table 2, while both analyses returned similar overall clustering metrics, segNASTIC as expected returned smaller clusters.

**Table 2.**
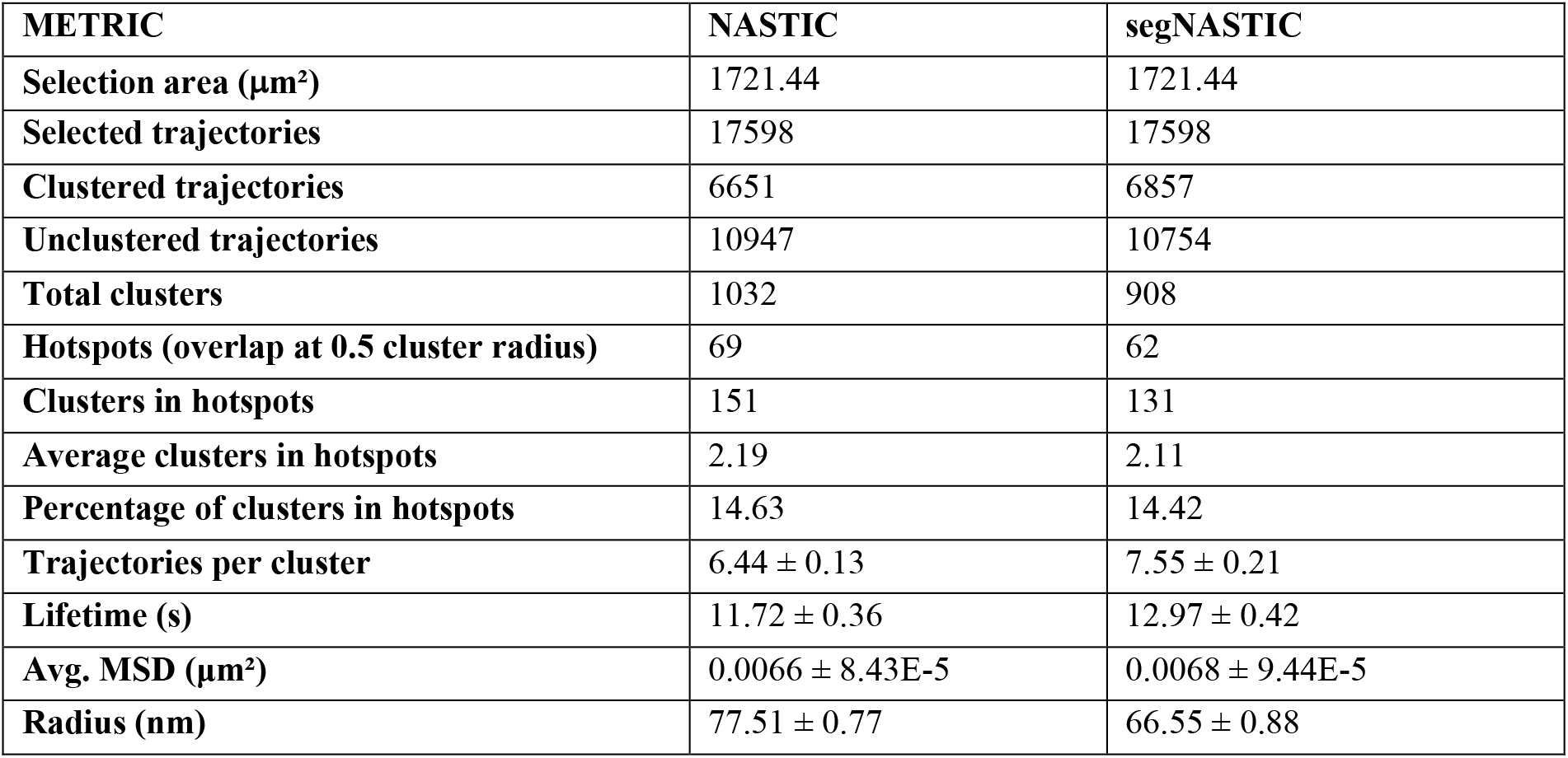
Comparison of NASTIC and segNASTIC on a typical dataset. For each of the clustering pipelines, the metrics returned from analysis of the same Sx1a-mEos2 dataset.

Given that different clustering algorithms use different mathematical approaches to determine molecular crossover, how does one truly define the spatial extent of a nanomolecular cluster? NASTIC uses the convex hull around all of the detections of the clustered trajectories, whilst segNASTIC uses the convex hull around the detections associated with overlapping trajectory segments (and thus reports smaller clusters). Beyond practical experimental concerns with data density, the choice of algorithm ultimately rests on its ability to detect experimentally and biologically driven changes in molecular clustering.

## CONCLUSION

The increasing sophistication of SMLM research has resulted in its application to studying fundamental biological processes such as exocytic neuronal communication. However, SMLM is a young and rapidly evolving field, with multiple approaches to data acquisition and data analysis. DBSCAN, Voronoï tessellation and R-tree spatial indexing are extremely robust, widely used and highly optimised algorithms that in the context of SMLM allow determination of nanomolecular clustering using fundamentally different approaches to arrive at similar conclusions regarding the geometry of molecular interaction. All frameworks based on these algorithms have their strengths and drawbacks, and are reliant on user metric guidance or empirical determination of optimal parameters. The fundamental difference of the approach detailed in this article is that temporal information is used to inform the algorithm such that spatiotemporal clusters are intrinsic to the analysis, rather than having to be derived subsequent to purely spatial clustering using approaches such as tcPALM (time correlated photoactivated light microscopy^9,43^). NASTIC is a robust pipeline that delivers a useful temporal dimension to SMLM analysis without dramatic increases in analysis time. The ability to resolve spatiotemporal clusters opens an unprecedented window into the dynamics of molecules on defined regions of the cell membrane. Spatiotemporal indexing can also potentially be expanded to multiple color SMLM analysis of different target proteins in the same cell^44^, where its ability to resolve hotspots may offer insights into cluster dependence, where clusters of one molecule may be dependent upon the pre-existing clusters of another molecule.

## MATERIALS AND METHODS

### PC12 cell culture, Transfection and Plating

Pheochromocytoma (PC12) cells were maintained in Dulbecco’s Modified Eagle Medium (DMEM, containing sodium pyruvate) (Thermo), foetal bovine serum (7.5%, Gibco) and horse serum (7.5%, Gibco), and 0.5% GlutaMax (Thermo) at 37°C an 5% CO2. Cells were transfected by Lipofectamine®LTX with Plus Reagent (Thermo) as per the manufacturers’ instructions with 2 ug of DNA or 1 ug of each plasmid when co-transfected. Cells were replated onto 0.1mg/ml Poly-D-lysine (Sigma) on 29mm No.1.5 glass-bottomed petri dishes (Cellvis) 24 h post-transfection, and imaged 48 h post-transfection. Live PC12 cells were imaged in isotonic condition in buffer A (145 mM NaCl, 5 mM KCl, 1.2 mM Na2HPO4, 10 mM D-glucose, and 20 mM Hepes, pH 7.4) at 37°C.

### Plasmids and fluorescent nanobodies

pmEos2-Munc18-1 (SNM) (Munc18-1-mEos2), pmEos2-N1 syntaxin 1a (Sx1a-mEos2), and pEGFP-N1 syntaxin 1a (Sx1a-EGFP) have been previously described^11^. Fluorescently labelled antibodies (Synatpic Systems, anti-GFP Atto647N tagged nanobodies, Cat#: GFP sdAb - FluoTag-Q - N0301-At647N-L) were reconstituted as per the manufacturer’s instructions, and utilized at 3.19 pg/μl in live uPAINT experiments.

### SMLM acquisition

PC12 cells transfected with Sx1a-mEos2 or Munc18-1-mEos2 were analysed by sptPALM. PC12 cells transfected with Sx1a-EGFP were analyzed by universal point accumulation in nanoscale topography (uPAINT) essentially as described in ^42^, and tracked using anti-GFP Atto647N tagged nanobodies ^40,45,46^ at 3.19 pg/µl.

Live transfected cells were visualized on a Roper Scientific Total Internal Reflection Fluoresecence (TIRF) microscope equipped with an iLas 2 double laser illuminator (Roper Scientific), a Nikon CFI Apo TIRF 100×/1.49 NA oil-immersion objective, and an Evolve 512 Delta EMCCD camera (Photometrics). Time-lapse TIRF movies (16,000 frames) were captured at 50 Hz for ∼320 s at 37 °C.

For single particle tracking photoactivated localization microscopy (sptPALM) analysis, samples were illuminated simultaneously with a 405 nm laser (Stradus, Vortan Laser Technology) to photoactivate mEos2-tagged proteins, and a 561 nm laser (Jive, Cobolt Lasers) for excitation of the photoconverted mEos2. A double beam splitter (LF488/561-A-000, Semrock) and a double-band emitter (FF01-523/610-25, Semrock) were used to isolate the mEos2 signal from autofluorescence and background signals. To achieve optimal spatial and temporal separation of stochaNASTIC mEos2 blinking, the power of the 405 nm and 561 nm laser was adjusted to 4mW and 22mW respectively at the focal plane.

For universal point accumulation for imaging in nanoscale topography (uPAINT), experiments were performed following Giannone et al. (2010). An anti-GFP nanobody (Kubala et al., 2010) tagged with ATTO 647N-NHS-ester (Atto-Tec) was used to track Sx1a-EGFP single molecules. PC 12 cells were double transfected with Munc18-1-mEos2 and Sx1a-EGFP to perform dual-color imaging. ATTO 647N–tagged anti-GFP nanobodies were added at a very low concentration for low level stochastic labelling. Time-lapse TIRF movies (16,000 frames) were recorded at 50 Hz at 37°C on TIRF microscope (Roper Technologies) equipped with an ILas2 double-laser illuminator (Roper Technologies). Each cell was imaged in both control condition and stimulated condition (2 mM BaCl2). A quadruple beam splitter (LF 405/488/561/635-A-000-ZHE; Semrock) and a quad band emitter (FF01-446/510/581/703-25; Semrock) were used for illumination. The power of the 635-nm laser used was 60% of initial laser power (200 mW).

All SMLM data were acquired using MetaMorph (Meta-Morph Microscopy Automation and Image Analysis Software, version 7.7.8; Molecular Devices), and further processed using the PalmTracer software^47^. NASTIC uses trajectory data in the TRXYT format, consisting of 4 headerless space separated columns corresponding to trajectory number, x co-ordinate (µm), y co-ordinate (µm), detection time (s). In this study, PalmTracer output was converted to TRXYT format using a custom Matlab script. Typical TRXYT data:

~~~
1 9.0117 39.86 0.02
1 8.9603 39.837 0.04
1 9.093 39.958 0.06
1 9.0645 39.975 0.08
etc
~~~

### Software

While initial investigations were performed using multiple command line driven Python scripts, we subsequently consolidated the analysis into two single-script graphic user interfaces (GUIs) suitable for general use by non-programmers, titled ***NASTIC*** (**Na**noscale **S**patio **T**emporal **I**ndexing **C**lustering) and ***Segment NASTIC (segNASTIC)***, for the whole trajectory and segment versions respectively. The two programs, collectively referred to as ***NASTIC***, require Python 3.8 or greater, and a number of modules which can be installed using:

~~~
python -m pip install numpy matplotlib pysimplegui rtree scipy sklearn
~~~

The GUI was constructed using the ***tk*** version of ***PySimpleGui*** (https://pysimplegui.readthedocs.io/). ***NASTIC*** was developed under and should run without issues on 64 bit Windows 7 and Windows 10. It will run with minor ***tk*** interface anomalies under Linux, but the authors cannot guarantee optimum performance under MacOS. NASTIC uses the Python ***multiprocessing*** module to fork computationally intensive operations onto all available cores of the physical or virtual machine on which it runs. This results in a dramatic increase in performance, but precludes packaging into standalone executables using ***PyInstaller***, and ***NASTIC*** will therefore only run on a computer with a Python 3.8+ environment.

The ***NASTIC*** interface divides the analysis workflow into a series of tabs which allow the user to: (1) load, screen and display the detections from an SMLM TRXYT file; (2) select one or more rectangular or irregular regions of interest (ROIs) and optionally adjust the density of selected trajectories within the ROIs; (3) adjust clustering parameters and apply NASTIC to the selected trajectories; (4) display and save the results of the clustering with a high degree of control over the presentation of trajectories, centroids and clusters; and (5) run a series of post clustering analyses to visualize metrics such as segment overlap, MSD, diffusion coefficient and cluster overlap probabilities, and save tabulated data for downstream comparative analyses. Typical visualizations are shown in Fig S5. Visualized data are saved as 300 dot per inch (dpi) PNG files, and the main clustering images can be additionally saved at a user specified dpi in a range of user specified file formats (EPS, PDF, PNG, PS, SVG, TIF). The co-ordinates of the ROIs are saved as tab separated ***roi_coordinates***.***tsv***: ROI, x co-ordinate (µm), y co-ordinate (µm) which can be loaded back into the ***NASTIC*** ROI tab. Analysis metrics are saved as tab separated ***metrics***.***tsv***, which can be viewed in any spreadsheet or text editor, and further processed for comparative studies using ***NASTIC Wrangler*** or other software.

Comparative analysis of multiple tabulated data files (generated during step 5 above) from 2 experimental conditions was performed using another simple Python GUI designated ***NASTIC Wrangler***. This program allows the user to specify two directories, each consisting of further subdirectories containing saved tabulated metrics data. ***NASTIC Wrangler*** recursively reads and compiles the tabulated metrics data from the subdirectories of each specified directory, and outputs a series of annotated comparative bar plots, thereby allowing the user to examine the degree and significance of change of a number of spatiotemporal clustering metrics. The significance of any differences in bar plots between conditions is determined by unpaired Student’s *t*-test.

## Code availability

All Python code is available at https://github.com/tristanwallis/smlm_clustering. ***NASTIC*** and associated scripts are released under a Creative Commons CC BY 4.0 licence: you are free to use and modify the code on the proviso that you make any changes freely available, acknowledge the original authors in derivative works and do not release said works under a more restrictive licence.

## AUTHOR CONTRIBUTIONS

F.A.M. and T.P.W. conceived the project, T.P.W implemented NASTIC in Python, H.H. assisted with code debugging and GUI implementation, A.J. performed super-resolution experiments under the supervision of M.J. and G.B., T.P.W. performed all clustering analyses with input from R.S.G., MSD analyses with input from N.D., and wrote the manuscript with input from the other authors.

## DATA AVAILABLITY

All processed data will be made available upon request and in The University of Queensland data repository, UQ eSpace.

## ACKNOWLEDGMENTS

We thank Rowan Tweedale for critical appraisal of the manuscript, the IT department at the Queensland Brain Institute (QBI), and Rumelo Amor and all past and present members of the Advanced Microscopy and Microanalysis Facility at QBI for their outstanding microscopy support. The work was supported by an Australian Research Council (ARC) Discovery Project grant (DP190100674), an ARC Linkage Infrastructure Equipment and Facilities grant (LE130100078) and a National Health and Medical Research Council (NHMRC) Senior Research Fellowship (GNT1155794) to FAM as well as an ARC Discovery Early Career Research Award (DE190100565) to MJ and NHMRC Project Grant (GNT 1147600) to ND.

## SUPPLEMENTARY FIGURES

**Figure S1.**
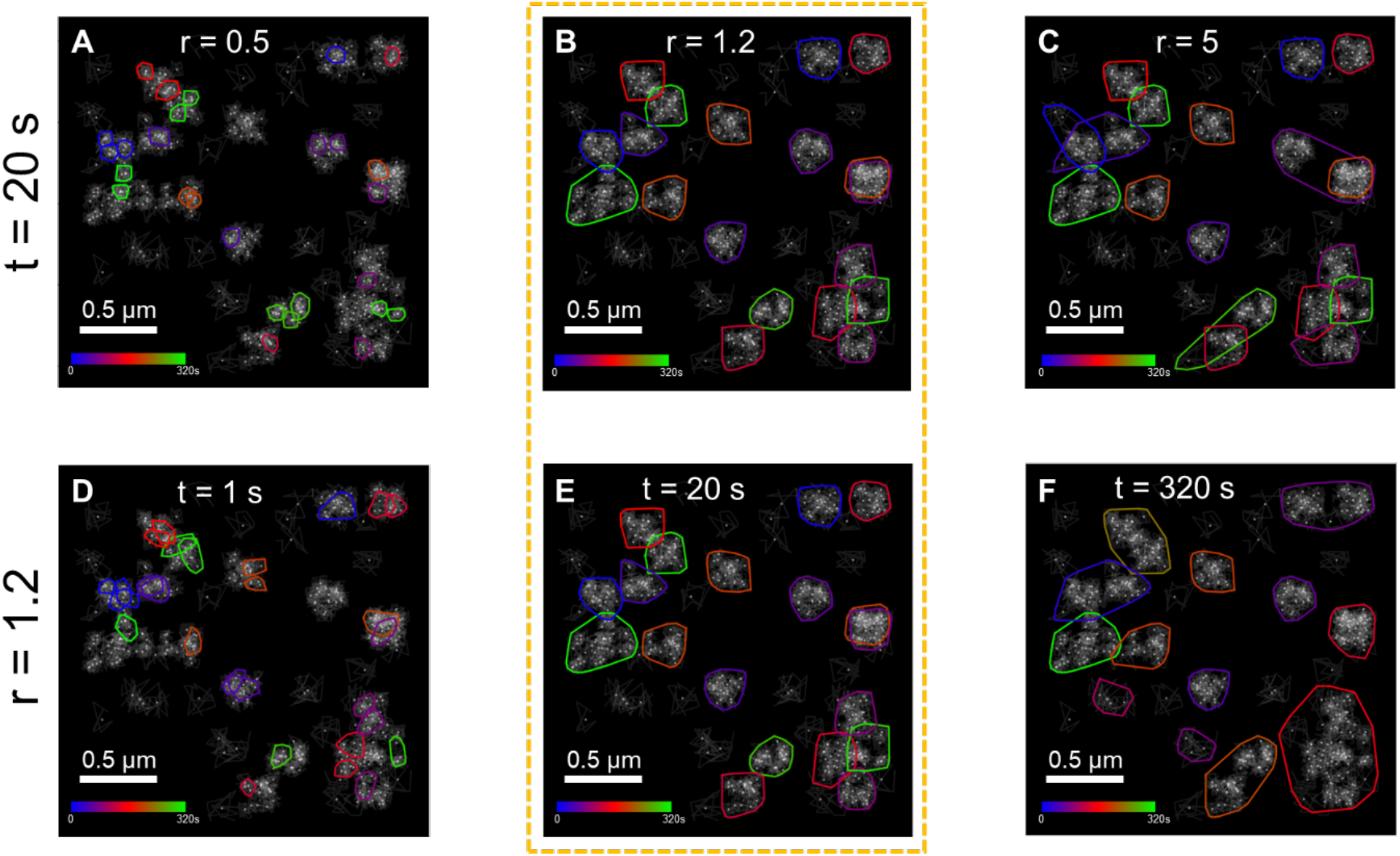
Bounding box radius and time window affect spatiotemporal clustering. *In silico* random walk trajectory data consisting of 20 equivalently sized spatiotemporal clusters where each cluster contains 20 trajectories distributed within 10 s. Clusters are randomly distributed within a 320 s “acquisition” window. The spatial centroid of each trajectory is represented as a dark dot. **Upper panels:** Spatiotemporal clustering was performed with a time window of 20 s, with each trajectory’s bounding box radius multiplied by the indicated factor r. **Lower panels:** Spatiotemporal clustering was performed with a bounding box radius factor of 1.2, and an indicated time window t. A cluster is defined as 3 or more proximal centroids. Cluster boundaries represent the extent of the detections associated with clustered trajectories, and are colored according to the average detection time as indicated by the color bar. Dotted box represents clustering obtained with r = 1.2, t= 20 s, most representative of the input data.

**Figure S2.**
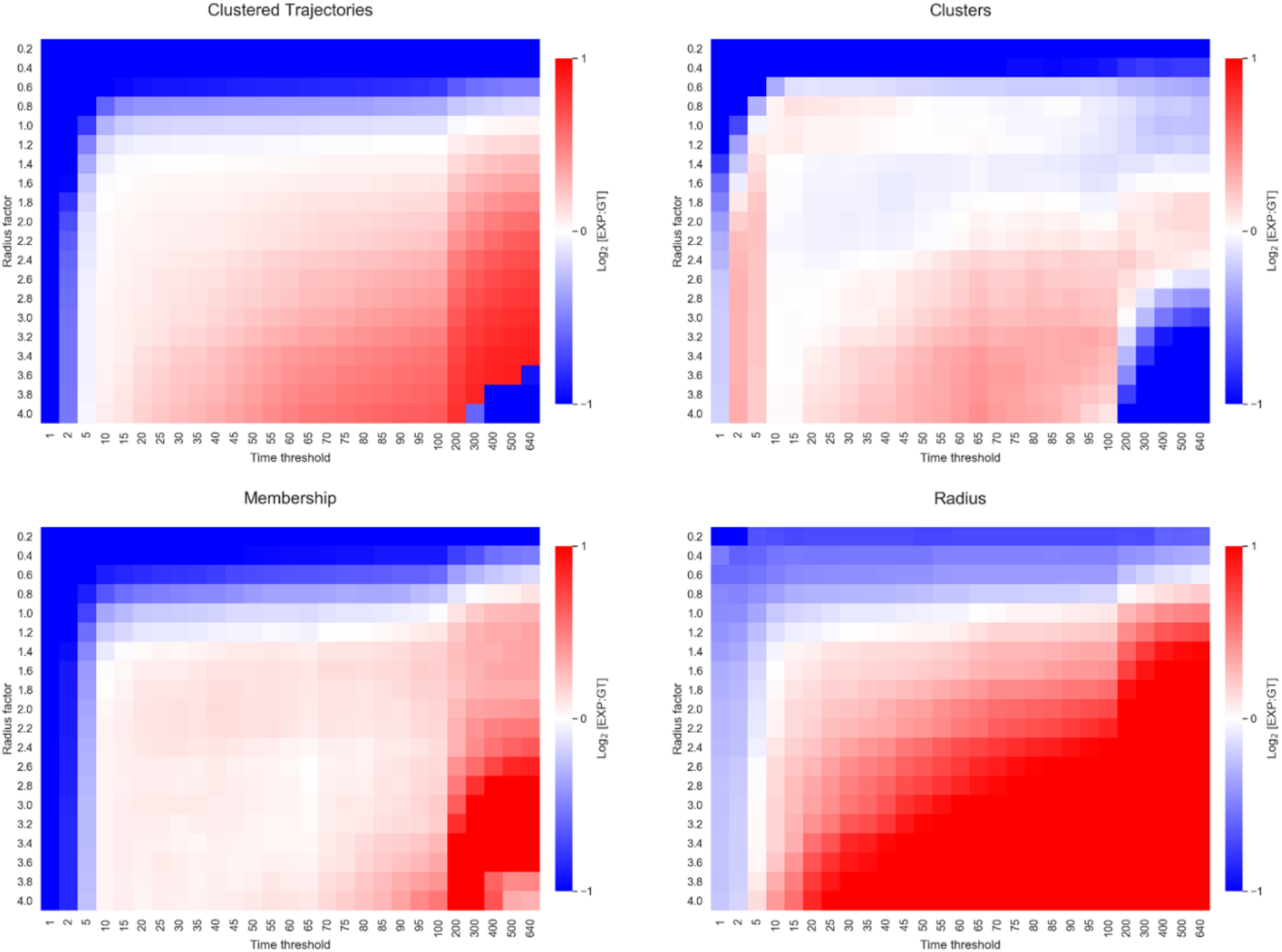
Effect of bounding box radius and time window on spatiotemporal clustering metrics. *In silico* random walk trajectory data consisting of 1095 trajectories in 140 spatiotemporally unique clusters with 7.82 ± 0.16 trajectories per cluster, with cluster radii of 74.86 ± 5.29 nm. The data also contained a background of 1000 randomly spatiotemporally distributed unclustered trajectories. Clusters are randomly distributed within a 320 s “acquisition” window. For a given metric, each pixel represents the log_2_ ratio of the experimental observed (EXP) value to the ground truth (GT). Ratios < -1 and > 1 are displayed as 1 and -1 respectively.

**Figure S3.**
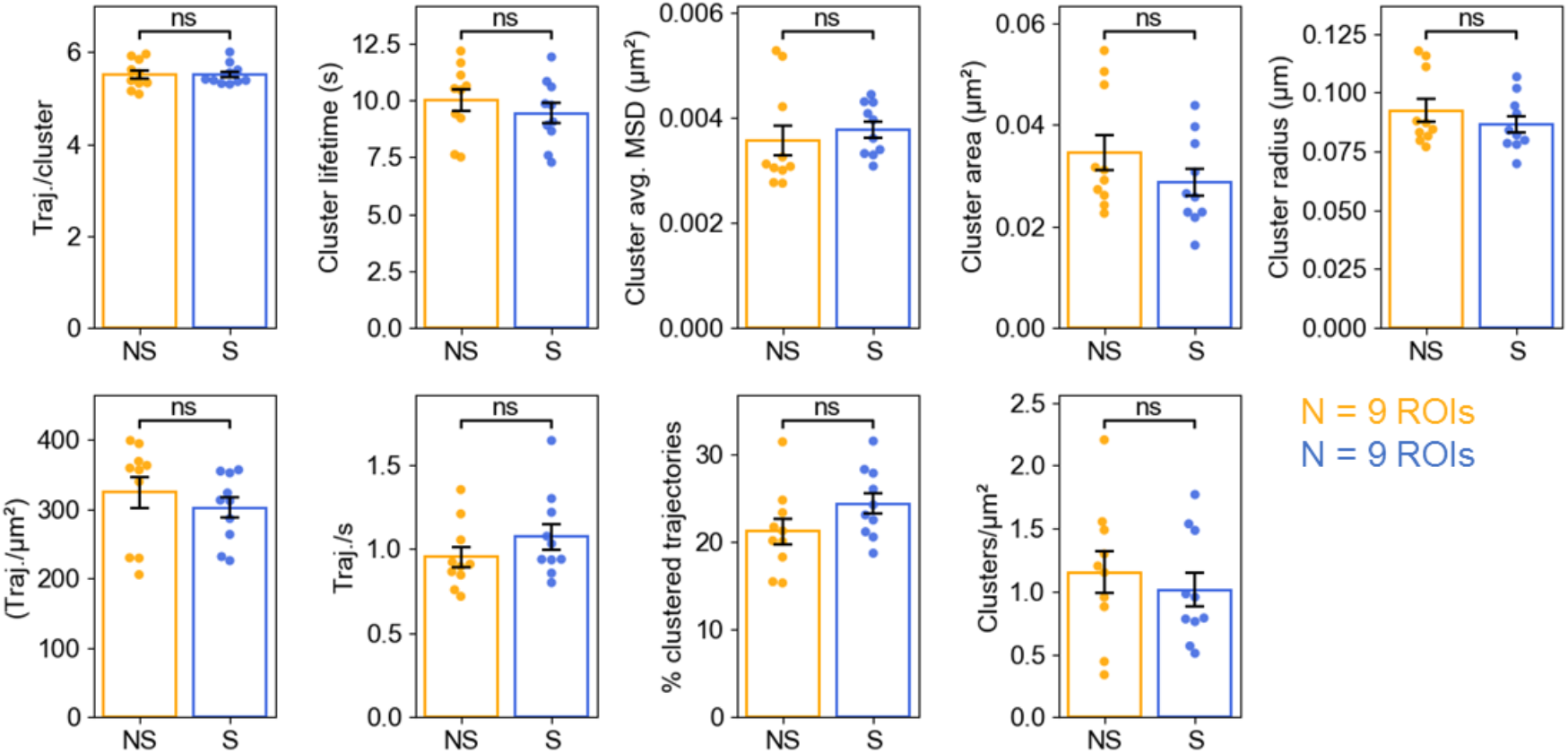
Difference in average cluster metrics in response to stimulated exocytosis. Munc18-1-mEos sptPALM data acquired from unstimulated PC12 cells (N=9) and PC12 cells stimulated with 2mM BaCl_2_ (N=9). (**A)** Representative image of clusters identified using spatiotemporal indexing with ***r*** = 1.2, ***t*** = 20 s. (**B)** Comparison of indicated average cluster metrics. The significance of the difference between conditions was determined by unpaired *t*-test (ns = no significance, *** = p < 0.001, **** = p < 0.0001). NS = no stimulation, S = stimulation with 2mM BaCl_2_

**Figure S4.**
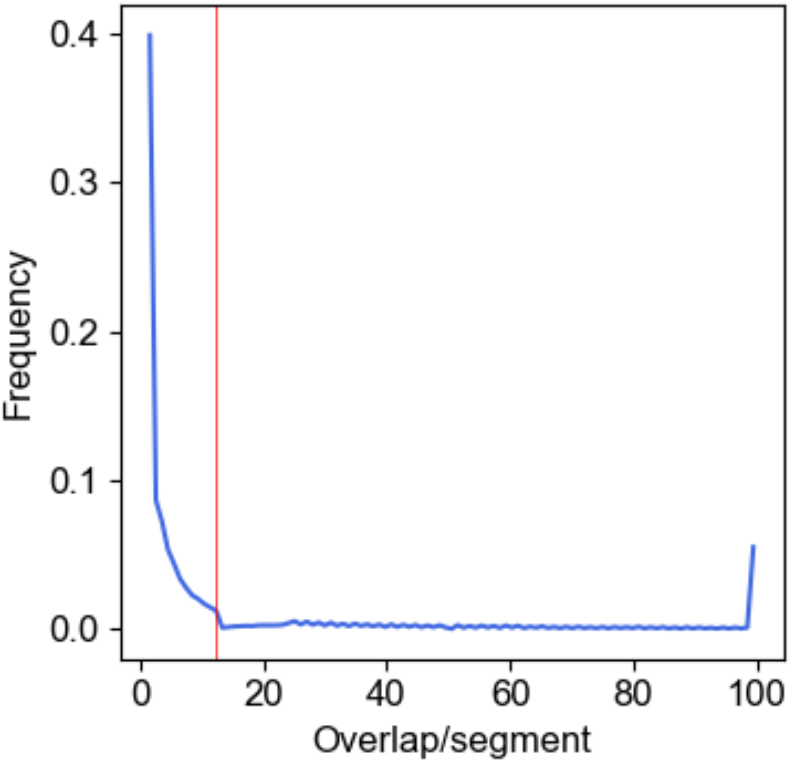
Distribution of segment overlap. Sx1a-GFP imaged by uPAINT using Atto-647 labelled anti-GFP nanobodies in PC12 cells. The vertical red line corresponds to the average segment overlap.

**Figure S5.**
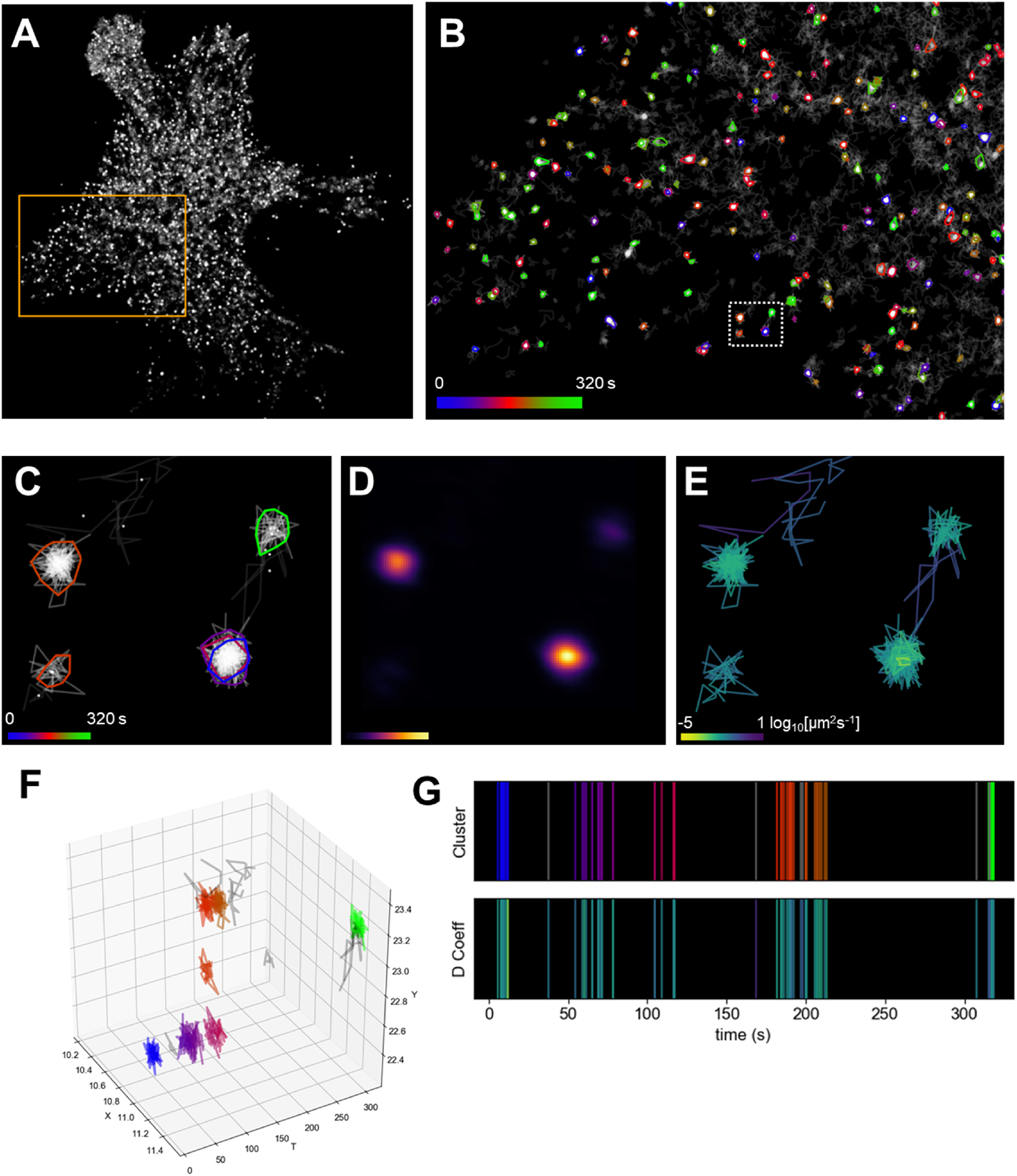
Typical NASTIC visualizations. NASTIC allows the user to evaluate nanomolecular clustering through a number of visualizations: (A) Raw acquisition data showing all molecular detections, with a region of interest (ROI) highlighted. (B) Spatiotemporal clustering of the selected trajectories within the ROI, with a region highlighted for enlargement. (C) Enlargement of highlighted area showing individual trajectories and their centroids, with clusters highlighted and color coded according to their time in the acquisition. (D) 2D Kernel density estimation of the detections associated with the selected trajectories, with brighter blobs corresponding to higher density. (E) Instantaneous diffusion coefficient, with each trajectory colored according to the gradient of the first 4 time points in its mean square displacement. (F) 3D plot of the selected trajectories, rotated to show the temporal separation of the clusters. (G) 1D plot of the selected trajectories where each vertical bar represents a single trajectory, colored according to its cluster status (top panel) or instantaneous diffusion coefficient (bottom panel).

## REFERENCES

1 Choquet, D., Sainlos, M. & Sibarita, J. B. Advanced imaging and labelling methods to decipher brain cell organization and function. Nat Rev Neurosci, (2021).

2 Kusumi, A., Tsunoyama, T. A., Hirosawa, K. M., Kasai, R. S. & Fujiwara, T. K. Tracking single molecules at work in living cells. Nat Chem Biol 10, 524–532, (2014).

3 Choquet, D. Linking Nanoscale Dynamics of AMPA Receptor Organization to Plasticity of Excitatory Synapses and Learning. J Neurosci 38, 9318–9329 (2018).

4 Goncalves, J. et al. Nanoscale co-organization and coactivation of AMPAR, NMDAR, and mGluR at excitatory synapses. Proc Natl Acad Sci U S A 117, 14503–14511, (2020).

5 Bademosi, A. T. et al. In Vivo Single-Molecule Tracking at the Drosophila Presynaptic Motor Nerve Terminal. J Vis Exp, (2018).

6 Bademosi, A. T. et al. In vivo single-molecule imaging of syntaxin1A reveals polyphosphoinositide- and activity-dependent trapping in presynaptic nanoclusters. Nat Commun 8, 13660, (2017).

7 Bademosi, A. T. et al. Trapping of Syntaxin1a in Presynaptic Nanoclusters by a Clinically Relevant General Anesthetic. Cell Rep 22, 427–440, (2018).

8 Chai, Y. J. et al. Munc18-1 is a molecular chaperone for alpha-synuclein, controlling its self-replicating aggregation. J Cell Biol 214, 705–718, (2016).

9 Gormal, R. S. et al. Modular transient nanoclustering of activated beta2-adrenergic receptors revealed by single-molecule tracking of conformation-specific nanobodies. Proc Natl Acad Sci U S A 117, 30476–30487, (2020).

10 Harper, C. B. et al. An Epilepsy-Associated SV2A Mutation Disrupts Synaptotagmin-1 Expression and Activity-Dependent Trafficking. J Neurosci 40, 4586-4595, d (2020).

11 Kasula, R. et al. The Munc18-1 domain 3a hinge-loop controls syntaxin-1A nanodomain assembly and engagement with the SNARE complex during secretory vesicle priming. J Cell Biol 214, 847–858, (2016).

12 Padmanabhan, P. et al. Need for speed: Super-resolving the dynamic nanoclustering of syntaxin-1 at exocytic fusion sites. Neuropharmacology 169, 107554, (2020).

13 Haas, K. T. et al. Pre-post synaptic alignment through neuroligin-1 tunes synaptic transmission efficiency. Elife 7, (2018).

14 Ester, M., Kriegel, H. P., Sander, J. & Xu, X. in KDD’96 Proceedings of the Second International Conference on Knowledge Discovery and Data Mining. 226–231.

15 Levet, F. et al. SR-Tesseler: a method to segment and quantify localization-based super-resolution microscopy data. Nat Methods 12, 1065-1071, doi:10.1038/nmeth.3579 (2015).

16 Andronov, L., Orlov, I., Lutz, Y., Vonesch, J. L. & Klaholz, B. P. ClusterViSu, a method for clustering of protein complexes by Voronoi tessellation in super-resolution microscopy. Sci Rep 6, 24084, (2016).

17 Khater, I. M., Nabi, I. R. & Hamarneh, G. A Review of Super-Resolution Single-Molecule Localization Microscopy Cluster Analysis and Quantification Methods. Patterns (N Y) 1, 100038, (2020).

18 Finkel, A. Quad trees, a data structure for retrieval on composite keys. Acta Informatica 4, 1–9 (1974).

19 Gutmann, A. R-Trees: A Dynamic Index Structure for Spatial Searching. Proceedings of the 1984 ACM SIGMOD international conference on Management of data – SIGMOD ‘84. p. 47 (1984).

20 Figueiredo, M. An R-tree Collision Detection Algorithm for Polygonal Models. Proceedings of the IASTED International Conference (2009).

21 Zhai, R. G. & Bellen, H. J. The architecture of the active zone in the presynaptic nerve terminal. Physiology (Bethesda) 19, 262–270, (2004).

22 Sudhof, T. C. The presynaptic active zone. Neuron 75, 11–25, (2012).

23 Tang, A. H. et al. A trans-synaptic nanocolumn aligns neurotransmitter release to receptors. Nature 536, 210–214, (2016).

24 Betzig, E. et al. Imaging intracellular fluorescent proteins at nanometer resolution. Science 313, 1642–1645, (2006).

25 Hess, S. T., Girirajan, T. P. & Mason, M. D. Ultra-high resolution imaging by fluorescence photoactivation localization microscopy. Biophys J 91, 4258–4272, (2006).

26 Manley, S. et al. High-density mapping of single-molecule trajectories with photoactivated localization microscopy. Nat Methods 5, 155-157, doi:10.1038/nmeth.1176 (2008).

27 Sudhof, T. C. & Rothman, J. E. Membrane fusion: grappling with SNARE and SM proteins. Science 323, 474–477, (2009).

28 Han, L. et al. Rescue of Munc18-1 and -2 double knockdown reveals the essential functions of interaction between Munc18 and closed syntaxin in PC12 cells. Mol Biol Cell 20, 4962–4975, (2009).

29 Rickman, C., Meunier, F. A., Binz, T. & Davletov, B. High affinity interaction of syntaxin and SNAP-25 on the plasma membrane is abolished by botulinum toxin E. J Biol Chem 279, 644–651, (2004).

30 Meunier, F. A. & Gutierrez, L. M. Captivating New Roles of F-Actin Cortex in Exocytosis and Bulk Endocytosis in Neurosecretory Cells. Trends Neurosci 39, 605–613, (2016).

31 Malintan, N. T. et al. Abrogating Munc18-1-SNARE complex interaction has limited impact on exocytosis in PC12 cells. J Biol Chem 284, 21637–21646, (2009).

32 Martin, S. et al. The Munc18-1 domain 3a loop is essential for neuroexocytosis but not for syntaxin-1A transport to the plasma membrane. J Cell Sci 126, 2353–2360, (2013).

33 Papadopulos, A. et al. Activity-driven relaxation of the cortical actomyosin II network synchronizes Munc18-1-dependent neurosecretory vesicle docking. Nat Commun 6, 6297, (2015).

34 Ullrich, A. et al. Dynamical Organization of Syntaxin-1A at the Presynaptic Active Zone. PLoS Comput Biol 11, e1004407, (2015).

35 Jahn, R. & Fasshauer, D. Molecular machines governing exocytosis of synaptic vesicles. Nature 490, 201–207, (2012).

36 Angelov, B. & Angelova, A. Nanoscale clustering of the neurotrophin receptor TrkB revealed by super-resolution STED microscopy. Nanoscale 9, 9797–9804, (2017).

37 Monnier, N. et al. Inferring transient particle transport dynamics in live cells. Nat Methods 12, 838–840, (2015).

38 Persson, F., Linden, M., Unoson, C. & Elf, J. Extracting intracellular diffusive states and transition rates from single-molecule tracking data. Nat Methods 10, 265–269, (2013).

39 Padmanabhan, P., Martinez-Marmol, R., Xia, D., Gotz, J. & Meunier, F. A. Frontotemporal dementia mutant Tau promotes aberrant Fyn nanoclustering in hippocampal dendritic spines. Elife 8, (2019).

40 Joensuu, M. et al. Subdiffractional tracking of internalized molecules reveals heterogeneous motion states of synaptic vesicles. J Cell Biol 215, 277–292, (2016).

41 Ripley, B. D. Modeling Spatial Patterns. J Roy Stat Soc B Met 39, 172–212 (1977).

42 Giannone, G., Hosy, E., Sibarita, J. B., Choquet, D. & Cognet, L. High-content super-resolution imaging of live cell by uPAINT. Methods Mol Biol 950, 95–110, (2013).

43 Cisse, II et al. Real-time dynamics of RNA polymerase II clustering in live human cells. Science 341, 664–667, (2013).

44 Subach, F. V. et al. Photoactivatable mCherry for high-resolution two-color fluorescence microscopy. Nat Methods 6, 153–159, (2009).

45 Kubala, M. H., Kovtun, O., Alexandrov, K. & Collins, B. M. Structural and thermodynamic analysis of the GFP:GFP-nanobody complex. Protein Sci 19, 2389–2401, (2010).

46 Joensuu, M. et al. Visualizing endocytic recycling and trafficking in live neurons by subdiffractional tracking of internalized molecules. Nat Protoc 12, 2590–2622, (2017).

47 Kechkar, A., Nair, D., Heilemann, M., Choquet, D. & Sibarita, J. B. Real-time analysis and visualization for single-molecule based super-resolution microscopy. PLoS One 8, e62918, (2013).

